# Prdm16-dependent antigen-presenting cells induce tolerance to intestinal antigens

**DOI:** 10.1101/2024.07.23.604803

**Authors:** Liuhui Fu, Rabi Upadhyay, Maria Pokrovskii, Francis M. Chen, Gabriela Romero-Meza, Adam Griesemer, Dan R. Littman

**Affiliations:** Department of Cell Biology, New York University School of Medicine, New York, NY, USA; Perlmutter Cancer Center, NYU Langone Health, New York, NY, USA; Calico Life Sciences, LLC, South San Francisco, CA, USA; NYU Langone Transplant Institute, NYU Langone Health, New York, NY, USA; Howard Hughes Medical Institute, New York, NY, USA

## Abstract

The gastrointestinal tract is continuously exposed to foreign antigens in food and commensal microbes with potential to induce adaptive immune responses. Peripherally induced T regulatory (pTreg) cells are essential for mitigating inflammatory responses to these agents^1–4^. While RORγt^+^ antigen-presenting cells (RORγt-APCs) were shown to program gut microbiota-specific pTreg^5–7^, their definition remains incomplete, and the APC responsible for food tolerance has remained elusive. Here, we identify a distinct subset of RORγt-APCs, designated tolerogenic dendritic cells (tDC), required for differentiation of both food- and microbiota-specific pTreg cells and for establishment of oral tolerance. tDC development and function require expression of the transcription factors Prdm16 and RORγt, as well as a unique *Rorc(t)* cis-regulatory element. Gene expression, chromatin accessibility, and surface marker analysis establish tDC as myeloid in origin, distinct from ILC3, and sharing epigenetic profiles with classical DC. Upon genetic perturbation of tDC, we observe a substantial increase in food antigen-specific T helper 2 (Th2) cells in lieu of pTreg, leading to compromised tolerance in mouse models of asthma and food allergy. Single-cell analyses of freshly resected mesenteric lymph nodes from a human organ donor, as well as multiple specimens of human intestine and tonsil, reveal candidate tDC with co-expression of *PRDM16* and *RORC* and an extensive transcriptome shared with mice, highlighting an evolutionarily conserved role across species. Our findings suggest that a better understanding of how tDC develop and how they regulate T cell responses to food and microbial antigens could offer new insights into developing therapeutic strategies for autoimmune and allergic diseases as well as organ transplant tolerance.

---

Antigen-presenting cells (APCs) play pivotal roles in orchestrating immune responses. They direct T cell outcomes, ranging from diverse effector to suppressive programs, according to potential threats from pathogenic microbes and the environmental context^8,9^. The adaptive immune response has evolved to be tolerant not only to self-antigens, but also to suppress responses to antigens present in the diet and in the mutualistic microbiota^10,11^. Tolerance to at least some commensal microbes with the potential to induce inflammation is mediated by pTregs that are programmed by recently-discovered RORγt-APCs^5–7^. Although these APCs express CD11c and Zbtb46, canonical markers of dendritic cells, they appear to have distinct features and have not been well characterized. They have been proposed to be MHCII^+^ type 3 innate lymphoid cells (ILC3) or subsets of recently described Janus cells (JC) and Thetis cells (TC), some of which express Aire^5–7^. Tolerance to dietary antigens is established through induction in the proximal alimentary tract of pTreg cells that restrain inflammatory responses to antigen encountered locally or distally. It is not known whether differentiation of such Treg cells is dependent on similar tolerogenic APCs. Classical dendritic cells (cDC) were proposed to be inducers of dietary antigen-specific pTregs, and it was also suggested that there is redundancy for this function among different types of APC^12,13^.

In this study, we set out to determine if the nuclear receptor RORγt, known to have critical roles in thymopoiesis, peripheral T cell differentiation, and development of innate lymphoid cells, is similarly required for the development and/or function of the newly described APC. We also investigated the role of RORγt-APCs in the induction of tolerance to oral antigen. We found that RORγt drives the development of a unique Prdm16-expressing, DC-like subset, with both RORγt and Prdm16 required for these APCs to induce pTregs that regulate inflammatory responses to food and microbiota antigens.

These APCs share many features with classical DCs, are present in mice across different ages, from neonatal to adult stages, and are also found in human tissues. We therefore suggest that they be designated as tolerogenic DC (tDC) that have critical roles in immune homeostasis and whose dysfunction likely contributes to multiple inflammatory and allergic diseases.

## Tolerogenic APC function requires *RORγt*

The novel tolerogenic APCs were discovered after genetic targeting by RORγt-cre of conditional alleles for MHCII, α_v_β_8_ integrin, and CCR7 resulted in loss of pTregs specific for microbiota antigens. While these studies indicated that the relevant APC expresses RORγt during ontogeny, it remained uncertain whether RORγt itself is required for these cells to develop or perform their function. We previously showed that inactivation of the same target genes in CD11c-cre mice also resulted in loss of pTreg-inducing APC function^5^.

To investigate a potential role for RORγt in these APCs, we therefore inactivated *Rorc(t)* in CD11c-cre mice and determined the fate of T cells specific for the large intestine pathobiont *Helicobacter hepaticus (Hh)*. Naïve *Hh*-specific CD4^+^ T cells from Hh7-2 TCR transgenic mice were transferred into *Hh*-colonized mice. Two weeks post-transfer, Hh7-2 T cells in the large intestine lamina propria (LILP) and mesenteric lymph nodes (mLN) of control mice displayed a predominance of pTregs expressing both RORγt and FOXP3 (Fig. 1a and Extended Data Fig. 1a,b). In contrast, in the LILP and mLN of *Cd11c*^cre^*Rorc(t)*^fl/gfp^ (*Rorc(t)*^Δ*CD11c*^) mice, the differentiation of adoptively transferred *Hh*-specific pTregs was abrogated. Instead, these mice exhibited an increase in RORγt- and T-bet-expressing *Hh*-specific T cells (Fig. 1a and Extended Data Fig. 1a,b), indicative of a shift towards a pro-inflammatory profile.

**Fig. 1.**
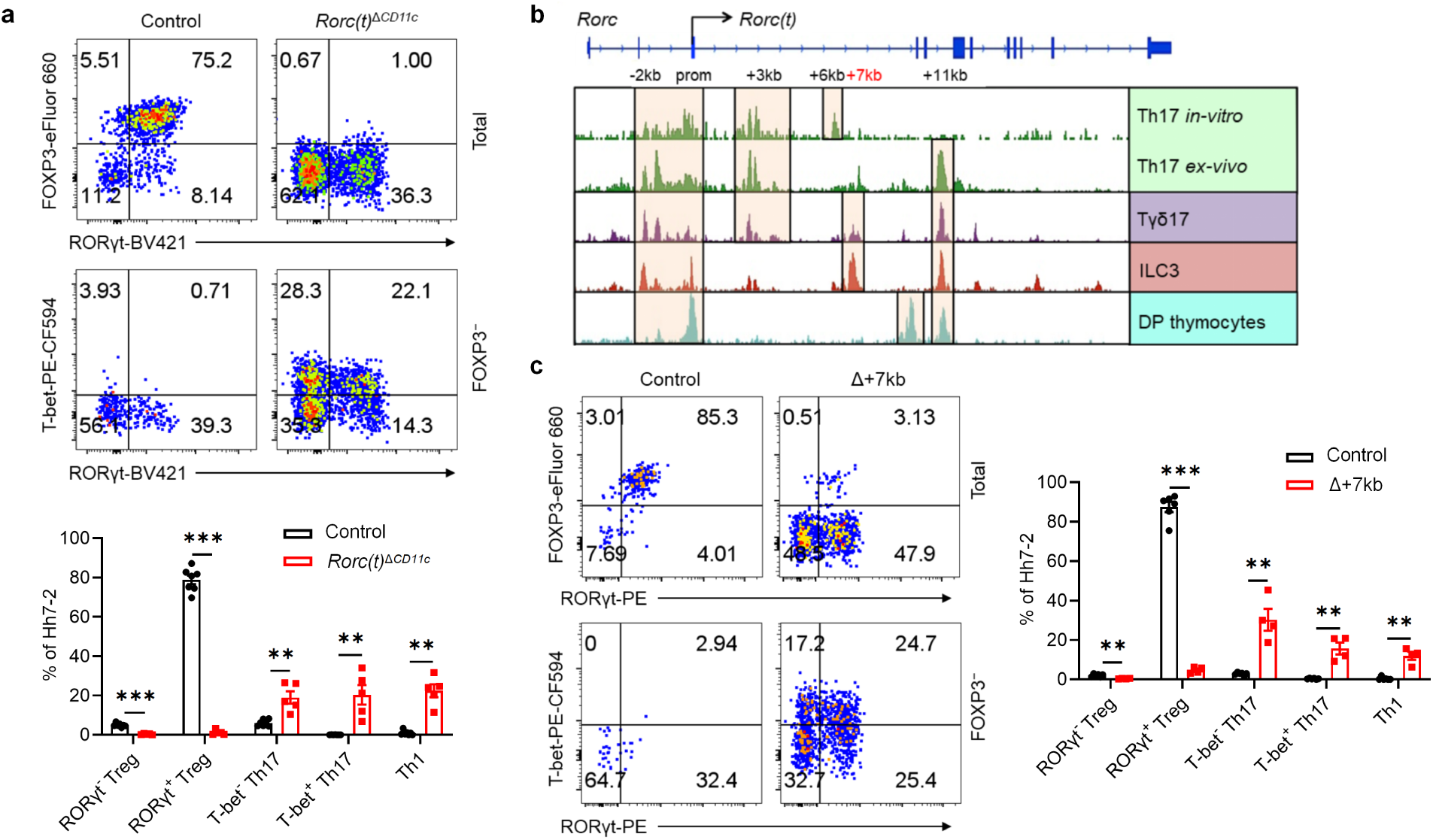
RORγt is required by tolerogenic APC to promote microbiota-specific pTreg differentiation. **a**, Representative flow cytometry plots (top) and frequencies (bottom) of *Hh*-specific pTreg (FOXP3^+^RORγt^+/−^), Th17 (FOXP3^−^RORγt^+^T-bet^+/−^) and Th1 (FOXP3^−^RORγt^−^T-bet^+^) cells in the LILP of *Hh*-colonized control (*Rorc(t)*^fl/gfp^, *Rorc(t)*^wt/gfp^, and *Cd11c*^cre^*Rorc(t)*^wt/gfp^; n = 7) and *Rorc(t)*^Δ*CD11c*^ (*Cd11c*^cre^*Rorc(t)*^fl/gfp^; n = 5) mice at 14 days after adoptive transfer of naïve Hh7-2tg CD4^+^ T cells. The upper panels of the flow cytometry plots are gated on total Hh7-2tg cells (CD45^+^B220^−^TCRγδ^−^TCRβ^+^CD4^+^Vβ6^+^CD90.1^+^), and the lower panels are gated on FOXP3^−^ Hh7-2tg cells. **b**, Bulk ATAC-seq data showing accessible regions in the *Rorc* locus of several RORγt-expressing cell types, including DP thymocytes (CD4^+^CD8^+^), *in vitro* differentiated Th17 cells, and SILP-derived Th17 (TCRβ^+^CD4^+^IL23R-GFP^+^) cells, Tγδ17 (TCRγδ^+^IL23R-GFP^+^) cells and ILC3 (Lin^−^IL-7R^+^Klrb1b^+^NK1.1^−^). **c**, Phenotype of *Hh*-specific T cells in the LILP of *Hh*-colonized control (*Rorc(t)* +7kb^+/+^; n = 6) and Δ+7kb (*Rorc(t)* +7kb^−/−^; n = 4) mice at 14 days after adoptive transfer of naïve Hh7-2tg CD4^+^ T cells. The flow cytometry plots are gated on total (upper) and FOXP3^−^ (lower) Hh7-2tg cells. Data in **a** are pooled from two independent experiments. Data in **c** are representative of two independent experiments. Data are means ± s.e.m.; ns, not significant; statistics were calculated by unpaired two-sided t-test.

Similarly, among endogenous T cells in these mutant mice there were fewer RORγt^+^ pTregs and substantially more inflammatory T helper 17 (Th17)/Th1 cells (Extended Data Fig. 1c). These results indicate that RORγt expression in CD11c lineage APCs is required for them to direct gut microbiota-specific pTreg cell differentiation.

## Lineage-specific *Rorc(t) cis* elements

RORγt is a transcription factor whose expression is largely confined to diverse lymphoid lineage cells, in which it contributes to distinct phenotypic programs^14–18^. Because different cis-regulatory elements (CREs) within a gene locus can govern its cell-specific expression, as best exemplified by the erythroid-specific enhancer in *Bcl11a^19^*, a therapeutic target for sickle cell disease, we hypothesized that distinct CREs within the *Rorc* locus may selectively modulate expression in the RORγt^+^ lineages, including the pTreg-inducing APCs. While prior research delineated several CREs involved in RORγt expression in Th17 cells and ILC3, *Rorc* regulatory regions in other cell types that express the transcription factor remain less well characterized^20–22^. To identify cell type-specific regulatory sequences, we conducted bulk ATAC-seq analyses in several RORγt-expressing cell types, including CD4^+^CD8^+^ thymocytes, *in vitro* differentiated Th17 cells, and small intestine lamina propria (SILP)-derived Th17 cells, Tγδ17 cells, and presumptive type 3 innate lymphoid cells (ILC3). These studies revealed distinct patterns of chromatin accessibility within the *Rorc* locus across the different cell types (Fig. 1b). Notably, regions situated +6kb and +7kb from the *Rorc(t)* transcription start site exhibited pronounced accessibility in *in vitro* differentiated Th17 cells and intestinal ILC3, respectively (Fig. 1b). Additionally, the +11kb element exhibited open chromatin configuration in all RORγt^+^ cell types, with the notable exception of *in vitro* polarized Th17 cells (Fig. 1b), consistent with our previous findings^21^. Further studies using dual reporter BAC transgenic mice specifically lacking a +3kb element (Tg (Δ+3kb *Rorc(t)*-mCherry);*Rorc(t)*^+/gfp^) indicated that *Rorc(t)* +3kb is a pivotal enhancer in Th17 and Tγδ17 cells *in vivo*, as well as *in vitro* differentiated Th17 cells, but not in ILC3 (Extended Data Fig. 2a,b).

To further explore the functional importance of the *Rorc(t)* +6 kb and +7kb elements, we engineered mice with deletions of these sequences. *Rorc(t)* +6kb^−/−^ (Δ+6kb) mice had significant reduction in both the proportion of Th17 cells in the SILP and level of RORγt expression within the remaining cells, but no change in ILC3 and Tγδ17 cells (Extended Data Fig. 2c,d). Notably, when naïve CD4 T cells from these knockout mice were subjected to Th17 differentiation conditions, they failed to upregulate RORγt (Extended Data Fig. 2f), confirming the regulatory importance of the +6kb region. In contrast, analysis of *Rorc(t)* +7kb^−/−^ (Δ+7kb) mice showed significantly reduced RORγt^+^ SILP ILC3 and Tγδ17 populations, with reduced RORγt expression in the remaining cells, but no effect in Th17 cells (Extended Data Fig. 2c,e). Naïve CD4^+^ T cells isolated from Δ+7kb mice displayed normal RORγt expression upon *in vitro* Th17 cell differentiation (Extended Data Fig. 2f), consistent with a role of this element only in innate-type lymphocytes.

The ILC3 subsets in the SILP and LILP of Δ+7kb mice were skewed towards Nkp46-expressing NCR^+^ ILC3, with reduction in CCR6-expressing LTi-like ILC3 (Extended Data Fig. 2g,h). Notably, the number of Peyer’s patches remained unchanged (Extended Data Fig. 2i), indicating that RORγt-dependent lymphoid tissue inducer (LTi) cells in mutant mice maintain sufficient functionality to support lymphoid organ development. When the Δ+7kb mice were challenged with the enteric pathogen *Citrobacter rodentium*, there was no difference in bacterial titers and weight loss compared to wild-type controls, nor was there any defect in IL-22 production (Extended Data Fig. 2j-l), which is essential for bacterial clearance^23^. These results suggested that mature ILC3 in the LILP of Δ+7kb mice retain functional capacity in response to *C. rodentium*, despite the reduction or loss of RORγt, which is required early for ILC3 development and for repression of T-bet, which, in turn, promotes expression of Nkp46 and transition to the ILC1 phenotype^24^. This finding is consistent with previous studies showing that while RORγt is crucial for restraining transcriptional networks associated with type 1 immunity, it is not essential for robust IL-22 production among mature ILC3^25,26^.

## Tolerogenic APC-specific *Rorc(t)* CRE

Despite the suggested maintenance of ILC3 function in the Δ+7kb mice, there was severe disruption in these animals of adoptively transferred *Hh*-specific pTreg cell differentiation in the LILP and mLN, accompanied by an expansion of inflammatory Th17/Th1 cells (Fig. 1c and Extended Data Fig. 3a,b). There was also a reduction of endogenous RORγt^+^ pTregs and increase of Th17/Th1 cells in the *Hh*-colonized mice (Extended Data Fig. 3c). Thus, the *Rorc(t)* +7kb CRE is required for RORγt-APCs to promote microbiota-specific pTreg cell differentiation even though known ILC3/LTi cell functions remain intact, which raises the possibility that such APCs belong to a different cell lineage than the ILCs.

Because RORγt is expressed in multiple immune system cell types, the T cell phenotype observed in the Δ+7kb mice could stem from direct or indirect effects. To clarify which cell types are affected by the deletion of the CRE, we established competitive bone marrow (BM) chimeric mice by co-transplanting CD45.1/2 wild-type BM along with either CD45.2 wild-type control or CD45.2 Δ+7kb BM into irradiated CD45.1 wild-type recipients (Extended Data Fig. 3d). We conducted a detailed analysis of donor chimerism across various intestinal immune cell subsets, including RORγt^+^ pTreg, ILC3, Th17, and Tγδ17 cells, normalizing these measurements to splenic B cells as an internal control. As expected, CCR6^+^RORγt^+^ ILC3 derived from the CD45.2 mutant donor were underrepresented, while Nkp46^+^RORγt^+^ ILC3 from the same donor were increased. However, RORγt^+^ pTreg, Th17 and Tγδ17 cells repopulated to similar extents in the mixed chimeras (Extended Data Fig. 3e,f). Analysis of RORγt levels within RORγt^+^ cell populations confirmed that that *Rorc(t)* +7kb intrinsically regulates RORγt expression in ILC3 and Tγδ17, but not in RORγt^+^ pTreg and Th17 cells (Extended Data Fig. 3g-i). Thus, the *Rorc(t)* +7kb mutation does not intrinsically affect T cell differentiation, which is consistent with the observed T cell phenotype being attributed to deficient RORγt-APC function.

## RORγt regulates Prdm16-expressing APCs

To better define the RORγt-dependent APCs required for intestinal pTreg cell differentiation, we performed single cell RNA sequencing (scRNA-seq) of MHCII-expressing innate immune system cells from mLN of control and mutant mice, including both Δ+7kb and *Rorc(t)*^Δ*CD11c*^ models. Cells harvested from Δ+7kb mutants and littermate controls at 3 weeks age yielded a total of 21,504 high-quality transcriptomes, comprising 11,150 and 10,354 cells from mutant and control animals, respectively (Fig. 2a). We analyzed these data with an unsupervised computation (Methods).

**Fig. 2.**
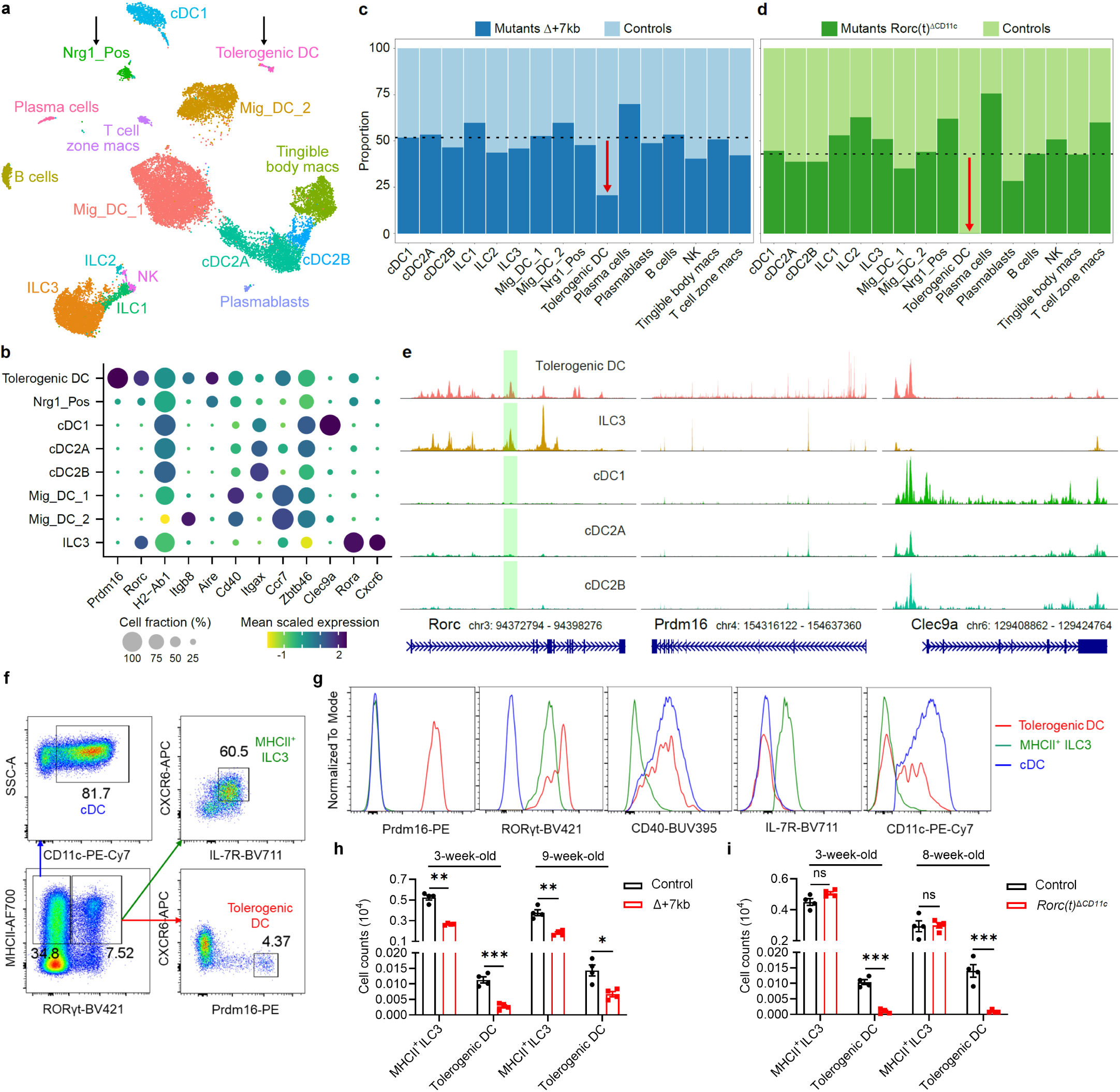
RORγt regulates development of Prdm16-expressing tolerogenic DC within mLN. **a**, UMAP representation of 21,504 transcriptomes obtained from scRNA-seq of MHCII-expressing innate immune system cells (CD45^+^Ly6G^−^B220^−^TCRγδ^−^TCRβ^−^MHCII^+^) in the mLN, combining data from 3-week-old Δ+7kb mutants (n = 7) and controls (n = 7) for joint clustering. Black arrows label two non-ILC3 clusters positive for *Rorc*. **b**, Dot plot of indicated clusters from **a** examining expression of genes previously ascribed to proposed RORγt-APC subsets. **c**, Stacked bar plots comparing proportion of each cluster in **a**, as derived from Δ+7kb mutant versus control animals. Dotted line at 51.2% indicates total contribution from Δ+7kb mutants. Red arrow indicates 60% loss within tolerogenic DC cluster of expected Δ+7kb contribution. **d**, Stacked bar plots analogous to experiment **a**-**c**, but now comparing proportion of cell clusters as derived from *Rorc(t)*^Δ*CD11c*^ mutants versus controls (n = 4 mice, each condition). Dotted line at 43% indicates total contribution from *Rorc(t)*^Δ*CD11c*^ mutants. Red arrow indicates complete loss within tolerogenic DC cluster of expected mutant contribution. **e**, Chromatin accessibility profiles for *Rorc*, *Prdm16*, and *Clec9a* loci across the indicated cell types. Green shaded range demarcates *Rorc(t)* +7kb CRE. **f**, Gating strategy for tolerogenic DC, MHCII^+^ ILC3 and classical DC (cDC). The bottom left flow cytometry plot is gated on CD45^+^Ly6G^−^B220^−^TCRγδ^−^TCRβ^−^ and was generated by concatenating samples from four wild-type mice. **g**, Expression of the indicated proteins in tolerogenic DC, MHCII^+^ ILC3 and cDC, as gated in **f**. **h**, Numbers of tolerogenic DC and MHCII^+^ ILC3 in the mLN of 3-week-old and 9-week-old control (n = 4) and Δ+7kb (n = 4) mice. **i**, Numbers of tolerogenic DC and MHCII^+^ ILC3 in the mLN of 3-week-old and 8-week-old control (n = 4) and *Rorc(t)*^Δ*CD11c*^ (n = 4) mice. Data in **f**-**i** are representative of two (**h**,**i**) or three (**f**,**g**) independent experiments. Data are means ± s.e.m.; ns, not significant; statistics were calculated by unpaired two-sided t-test.

Populations of DCs and ILCs and distinct clusters of B cell and macrophage lineages were readily annotated (Fig. 2b, Extended Data Fig. 4a). Juxtaposed ILCs were partitioned based on canonical expression of *Ncr1*/*Tbx21* (Type 1), *Il1rl1*/*Il4* (Type 2), or *Cxcr6/Ccr6/Nrp1* (Type 3), versus a contiguous population of natural killer cells positive for *Eomes* and *Gzma*. cDC1 occupied a distinct cluster positive for *Xcr1* and *Clec9a*. While subsets of cDC2 (all positive for *Sirpa*) were contiguous, they could be separated based on expression of *Clec4a4*/*Cd4*/*Dtx1* (cDC2A) versus *Cd209a*/*Cx3cr1*/*Mgl2* (cDC2B), with all *Tbx21* expression contained within the former, as previously reported^27^. We observed two sizable populations of *Ccr7*^+^ migratory DCs that were denoted Mig_DC_1 and Mig_DC_2, both consistent with a signature known to include *Socs2*, *Fas*, and *Cxcl16^5,28^*.

On querying for *Rorc* positive cells, beyond expected ILC3 there were two additional distinct clusters (arrows, Fig. 2a, Extended Data Fig. 4b). We denoted one as Nrg1_Pos, given exclusively high expression of neuregulin 1, a trophic factor known to coordinate various neuronal functions including axon guidance, myelination, and synapse formation (Extended Data Fig. 4c). Additional neuronal genes were also prominent (*Ncam1*, *Nrxn1*, *Nrn1*), as was expression of *Aire*, *Trp63*, and *Tnfrsf11b*/osteoprotegerin, in many ways compatible with the described TC I subset^6^. This cluster also harbored a signature closely associated with the fibroblastic reticular cell (FRC) subset found in mucosal lymph nodes^29^, including *Madcam1*, *Mfge8*, *Twist1*, *Vcam1*, and *Nid1*. Both non-ILC3 *Rorc*^+^ clusters expressed CD45, as well as MHCII at levels comparable to DC subsets. Therefore, Nrg1_Pos cells are not bona fide stromal cells nor medullary thymic epithelial cells (mTEC), but they harbor prominent neural, FRC, and mTEC attributes.

The second *Rorc*^+^ cluster displayed exclusively high expression of *Prdm16* (Fig. 2b, Extended Data Fig. 4d), previously shown to be variably expressed across TC subsets^6^. This singular appearance of *Prdm16* in our dataset seemed notable, given that it is a zinc finger protein and histone methyltransferase previously identified as a transcriptional regulator in multiple cell types, including brown adipocytes, cortical neurons, and vascular endothelial cells^30^. The *Prdm16*^+^ cluster was notably higher in *Itgb8*, *Aire*, *Cd40*, and *Ccr7* expression when compared to Nrg1_Pos cells (Fig. 2b).

After annotating, we deconvolved each cluster according to its origin from Δ+7kb mutant or control animals. While the contribution from each condition varied minimally around the numeric split of total high-quality transcriptomes captured (Fig. 2c, dotted line), we observed a striking 60% reduction of *Prdm16*^+^ cluster cells from the expected Δ+7kb contribution. Violin plots examining *Rorc* and *Ccr6* illustrate clear decreases of those gene expressions within the mutant-origin ILC3 population (Extended Data Fig. 4e), yet the overall cluster of ILC3 from mutant mice remained otherwise numerically intact. As there was also no apparent difference in the Nrg1_Pos distribution when comparing control versus Δ+7kb mutant mice, the results suggested that the specialized ability to induce pTreg is contained in the *Prdm16*^+^ population.

Comparison of mLN single cell transcriptomes from *Rorc(t)*^Δ*CD11c*^ (8,474 cells) and littermate control mice (11,245 cells) yielded results congruent with those observed in the Δ+7kb model. In these mutants, however, there was complete loss of the *Prdm16*^+^ cluster when we deconvolved biological conditions, despite a 43% overall contribution of total transcriptomes from *Rorc(t)*^Δ*CD11c*^ mice (Fig. 2d, dotted line). Together, the results are most consistent with a requirement for RORγt expression during development of Prdm16^+^ APCs and suggest that these cells are likely candidates for tolerogenic APCs in response to the microbiota.

## Characterization of Prdm16^+^ APCs as tDC

To further define the Prdm16^+^ APCs, we employed single cell assay for transposase-accessible chromatin using sequencing paired with transcriptomic readout (multiome scRNA-seq/scATAC-seq), resulting in 14,917 high quality nuclei from mLN of *Rorc(t)* fate-mapped mice. The results were computationally integrated^31^ with data from all prior control and mutant animals, generating the exact same populations as above (Extended Data Fig. 4f). A previous investigation had employed a similar multi-ome analysis, albeit on nuclei enriched via a *Rorc*-reporter animal^6^. We additionally integrated that raw data alongside our own, which proved to be a critical step before shared nearest neighbor clustering, as this provided the context of all other MHC II-expressing APCs found within mLN beyond *Rorc*-enriched cells, which were still observed in the three distinct clusters, ILC3, Nrg1_Pos and Prdm16^+^, suggesting that our strategy for computation and annotation was robust (Extended Data Fig. 4g).

Consistent with the gene expression data, scATAC-Seq revealed chromatin accessibility at both *Rorc* and *Prdm16* loci in *Prdm16*^+^ APCs (Fig. 2e). Close inspection of the *Rorc(t)* +7kb CRE (green shaded range), demonstrated prominent peaks in both *Prdm16*^+^ APCs and ILC3, as expected. Comparing chromatin accessibility of *Prdm16*^+^ APCs to that of ILC3, cDC1, cDC2A, and cDC2B revealed a consistent epigenetic pattern. For instance, although *Prdm16*^+^ APCs did not transcribe *Clec9a*, the locus was accessible, with conspicuous peaks noticeably shared across cDC populations but absent in ILC3. This pattern of chromatin landscape shared by *Prdm16*^+^ APCs and cDCs, but not ILC3, was observed at multiple loci, including *Csf1r*, *Flt3*, *Clecl0a*, *Sirpa*, *Itgax*, *Itgam*, and *Itgae* (Extended Data Fig. 5). We observed the converse for *Cxcr6*, with chromatin accessibility and gene expression in ILC3, but not in *Prdm16*^+^ APCs and cDCs.

Using a flow cytometry gating strategy for Prdm16^+^ APCs, MHCII^+^ ILC3 and cDC, we confirmed that Prdm16^+^ APCs co-expressed Prdm16 and RORγt at high levels (Fig. 2f,g). Prdm16^+^ APCs also expressed CD40 at levels comparable to cDC, supporting their involvement in T cell priming. In contrast to MHCII^+^ ILC3, Prdm16^+^ APCs and cDC lacked expression of IL-7R, CXCR6, and CD90 (Fig. 2g and Extended Data Fig. 6a). Notably, CD11c and CD11b expression was limited to a portion of Prdm16^+^ APCs, suggesting that there is partial downregulation of CD11c during lineage progression. In terms of side scatter (SSC), Prdm16^+^ APCs resembled cDC, while their forward scatter (FSC) was intermediate between MHCII^+^ ILC3 and cDC (Extended Data Fig. 6b). These results indicate that Prdm16^+^ APCs and MHCII^+^ ILC3 are distinct populations, with Prdm16^+^ APCs sharing more similarities with cDC. In addition, our analysis of Prdm16^+^ APC abundance across a developmental time frame, from 1-week-old to 12-week-old mice, revealed a slight increase in their numbers with age, followed by stabilization in adulthood (Extended Data Fig. 6c). This pattern aligns with the observed ability of adult mice to induce tolerance to gut microbiota^5,32^, suggesting that these APCs maintain functional persistence beyond early development.

Further validation by flow cytometry analysis revealed that, although the numbers and RORγt MFI of Prdm16^+^ APCs and MHCII^+^ ILC3 were decreased in the mLN of Δ+7kb mice, MHCII^+^ ILC3 number remained unchanged in *Rorc(t)*^Δ*CD11c*^ mice, whereas Prdm16^+^ APCs were depleted (Fig. 2h,i and Extended Data Fig. 6d,e), indicating that Prdm16^+^ APCs but not ILC3 are the potential pTreg-inducing APCs. To further test this hypothesis, we conditionally inactivated Prdm16 in RORγt-expressing cells (*Prdm16*^Δ*RORγt*^ mice) and assessed the fate of *Hh*-specific T cells. In these mice, MHCII^+^ ILC3 numbers were unchanged in the mLN and intestines, while Prdm16^+^ APCs were barely detectable when identified using Prdm16 as a marker (Extended Data Fig. 7a,b). In accordance, the differentiation of adoptively transferred *Hh*-specific pTreg was abolished in the LILP and mLN of *Hh*-colonized *Prdm16*^Δ*RORγt*^ mice, and was accompanied by an increase in inflammatory Th17/Th1 cells (Fig. 3a and Extended Data Fig. 7c,d), suggesting at least a loss of tolerogenic function in Prdm16^+^ APCs. Collectively, these findings establish Prdm16^+^ APCs as a distinct population closely resembling cDC, yet uniquely capable of inducing microbiota-specific pTregs, supporting their designation as tolerogenic dendritic cells (tDC).

**Fig. 3.**
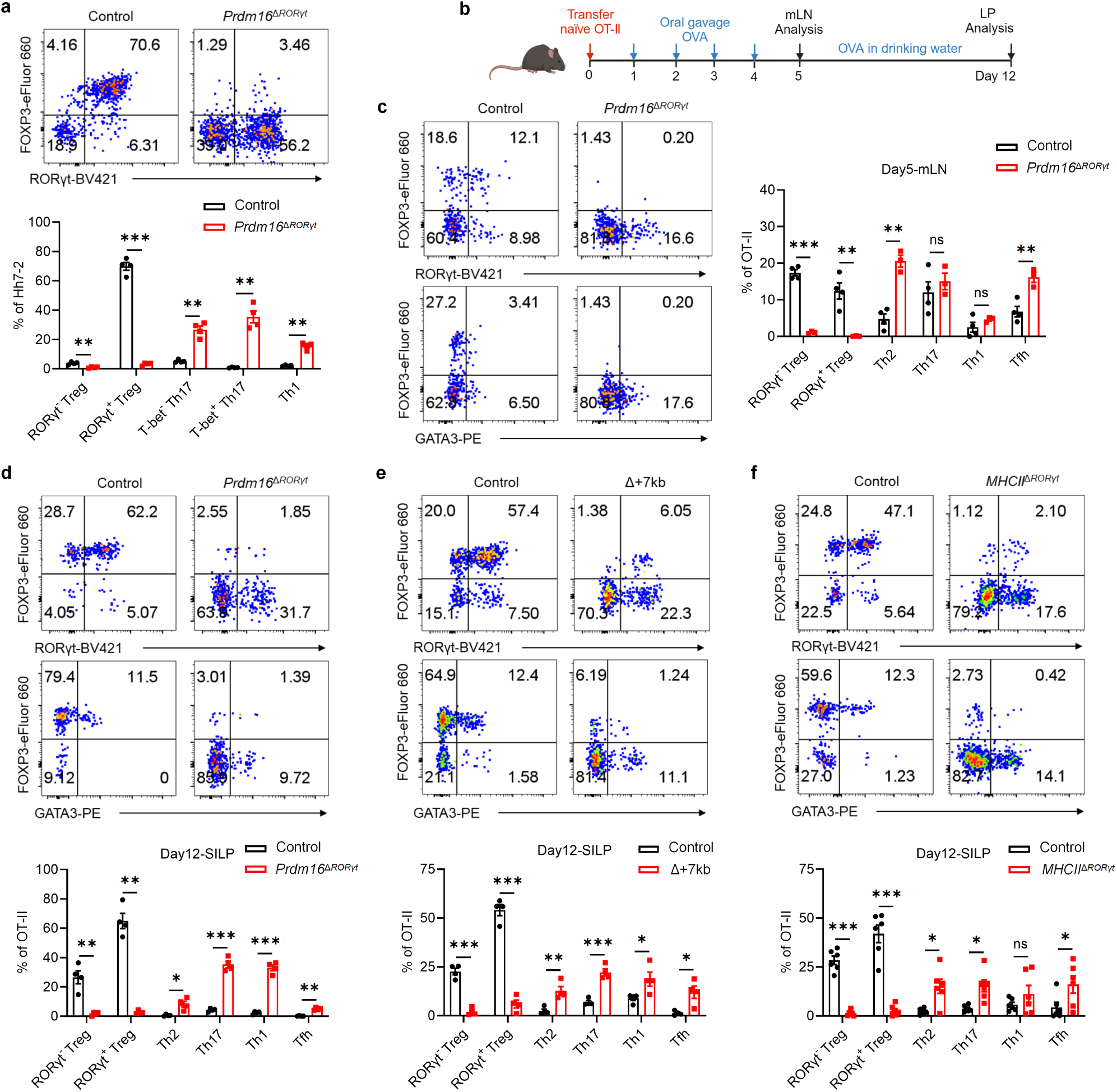
Prdm16-dependent tolerogenic DC promote both microbiota and food antigen-specific pTreg differentiation. **a**, Phenotype of *Hh*-specific T cells in the LILP of *Hh*-colonized control (*Prdm16*^fl/fl^, *Prdm16*^fl/wt^; n = 4) and *Prdm16*^Δ*RORγt*^ (*Rorc(t)*^cre^*Prdm16*^fl/fl^; n = 4) mice at 14 days after adoptive transfer of naïve Hh7-2tg CD4^+^ T cells. **b**, Experimental design for the experiments in **c**-**f**. OVA, ovalbumin; mLN, mesenteric lymph nodes; LP, lamina propria. **c**-**f**, Representative flow cytometry plots and frequencies of OT-II pTreg (FOXP3^+^RORγt^+/−^), Th2 (FOXP3^−^GATA3^+^), Th17 (FOXP3^−^GATA3^−^RORγt^+^), Th1 (FOXP3^−^GATA3^−^RORγt^−^T-bet^+^) and Tfh (FOXP3^−^GATA3^−^RORγt^−^T-bet^−^BCL6^+^) cells in the mLN (**c**) and SILP (**d**-**f**) of OVA-treated control and mutant mice at 5 and 12 days post-adoptive transfer of naïve OT-II CD4^+^ T cells. The flow cytometry plots are gated on total OT-II cells (CD45^+^B220^−^TCRγδ^−^TCRβ^+^CD4^+^Vα2^+^Vβ5.1/5.2^+^GFP^+^). **c**, mLN: control mice, n = 4; *Prdm16*^Δ*RORγt*^ mice, n = 3. **d**, SILP: control mice, n = 4; *Prdm16*^Δ*RORγt*^ mice, n = 4. **e**, SILP: control mice, n = 4; Δ+7kb mice, n = 4. **f**, SILP: control (*I-AB*^fl/fl^) mice, n = 6; *MHCII*^Δ*RORγt*^ (*Rorc(t)*^cre^*I-AB*^fl/fl^) mice, n = 6. Data in **f** are pooled from two independent experiments. Data in **a**,**d**,**e** are representative of two (**a**,**d**) or three (**e**) independent experiments. Data are means ± s.e.m.; ns, not significant; statistics were calculated by unpaired two-sided t-test.

## tDC required for oral tolerance

When Δ+7kb, *Rorc(t)*^Δ*CD11c*^ and *Prdm16*^Δ*RORγt*^ mice were examined in the absence of *H. hepaticus* colonization, we observed the expected reduction, compared to control mice, of RORγt^+^ pTregs within both SILP and LILP (Extended Data Fig. 7e,g,h). Intriguingly, this decrease did not coincide with an increase in Th17 cell proportions but was associated with a substantial elevation of intestinal Th2 cell levels (Extended Data Fig. 7e-h). Notably, in 40-week-old Δ+7kb mice, no overt intestinal inflammation was observed by hematoxylin and eosin (HE) staining. Nevertheless, these mice displayed features indicative of spontaneous type 2 gastrointestinal pathology^33,34^, including muscularis propria hypertrophy, increased small intestine length, and elevated serum total IgE levels (Extended Data Fig. 7i-l). These observations in the mutant mice suggested that tDC induce pTregs specific not only for microbiota, but also for food antigens with potential to induce allergy-related Th2 cells. To test this hypothesis, we transferred naïve ovalbumin (OVA)-specific CD4^+^ T cells from OT-II TCR transgenic mice into both control and mutant mice deficient for microbiota-specific pTreg induction, and administered OVA either via gavage or in drinking water (Fig. 3b). Interestingly, OVA-specific OT-II pTregs in wild-type mice consisted of both RORγt^-^ and RORγt^+^ phenotypes (Fig. 3c-f). At five days post-transfer, there were few OT-II pTregs in the mLN of *Prdm16*^Δ*RORγt*^, Δ+7kb and *Rorc(t)*^Δ*CD11c*^ mice, and, instead, the OT-II T cells displayed Th2 and T follicular helper (Tfh) phenotypes, with no notable changes in Th17 and Th1 profiles (Fig. 3c and Extended Data Fig. 8a-c). By the twelfth day post-transfer, the reduction in OT-II pTregs persisted in the intestines of *Prdm16*^Δ*RORγt*^, Δ+7kb and *MHCII*^Δ*RORγt*^ (*Rorc(t)*^cre^*I-AB*^fl/fl^) mice, coinciding with variable increases across all OT-II Th cell subsets (Fig. 3d-f and Extended Data Fig. 8d-f). These results highlight the critical role of tDC in promoting food antigen-specific pTreg cell differentiation. Dysfunction of these APCs correlates with intensified effector Th cell responses, although the specific Th cell subset favored depends on the local tissue environment.

Peripheral pTreg cells are pivotal in the induction and maintenance of oral tolerance, a critical mechanism that suppresses diverse immune responses not only in the gastrointestinal tract but also systemically^35^. We therefore aimed to explore whether tDC are essential for directing food antigen-specific pTregs to mediate oral tolerance. We first used an allergic lung response model, in which oral administration of antigen prior to sensitization results in pTreg-mediated inhibition of the inflammatory process. Mice were not pre-treated or were administered OVA intragastrically before being primed with OVA in alum and subsequently exposed to intranasal OVA challenge (Fig. 4a). Wild-type mice that had previously been fed OVA exhibited significant resistance to allergic lung inflammation, demonstrated by lower lung inflammation score, diminished eosinophil numbers in the bronchoalveolar lavage fluid (BALF) and lungs, reduced lung Th2 cells, and decreased levels of serum OVA-specific IgE and IgG1 compared to non-tolerized controls (Fig. 4b-d and Extended Data Fig. 9a-f). However, the same pre-feeding strategy failed to induce oral tolerance in Δ+7kb mice. These knockout mice showed similar increases in lung inflammation score, eosinophils, Th2 cells, and OVA-specific IgE and IgG1 production as non-tolerized Δ+7kb mice (Fig. 4b-d and Extended Data Fig. 9a-f). We then focused on OVA-specific T cell responses in these mice, using OVA:I-A^b^ tetramers to identify those cells. In tolerized wild-type mice, tetramer-positive T cells were significantly fewer compared to non-tolerized animals, and most cells were GATA3^+^FOXP3^+^ Treg cells, with limited RORγt expression (Fig. 4e,f and Extended Data Fig. 9g,h). In contrast, in both tolerized and non-tolerized Δ+7kb mice there was loss of OVA:I-A^b^ tetramer-binding pTregs in the lung, accompanied by an increase in tetramer-positive Th2, Th17, and Th1 cells, with a predominant increase in Th2 cells (Fig. 4e,f and Extended Data Fig. 9g-h), consistent with loss of tolerance. Similarly, OVA-pre-fed *Prdm16*^Δ*RORγt*^, *Rorc(t)*^Δ*CD11c*^ and *MHCII*^Δ*RORγt*^ mice exhibited comparable increases in lung eosinophils and Th2 cells to those observed in non-tolerized mutant mice (Fig. 4g-i and Extended Data Fig. 9i-k).

**Fig. 4.**
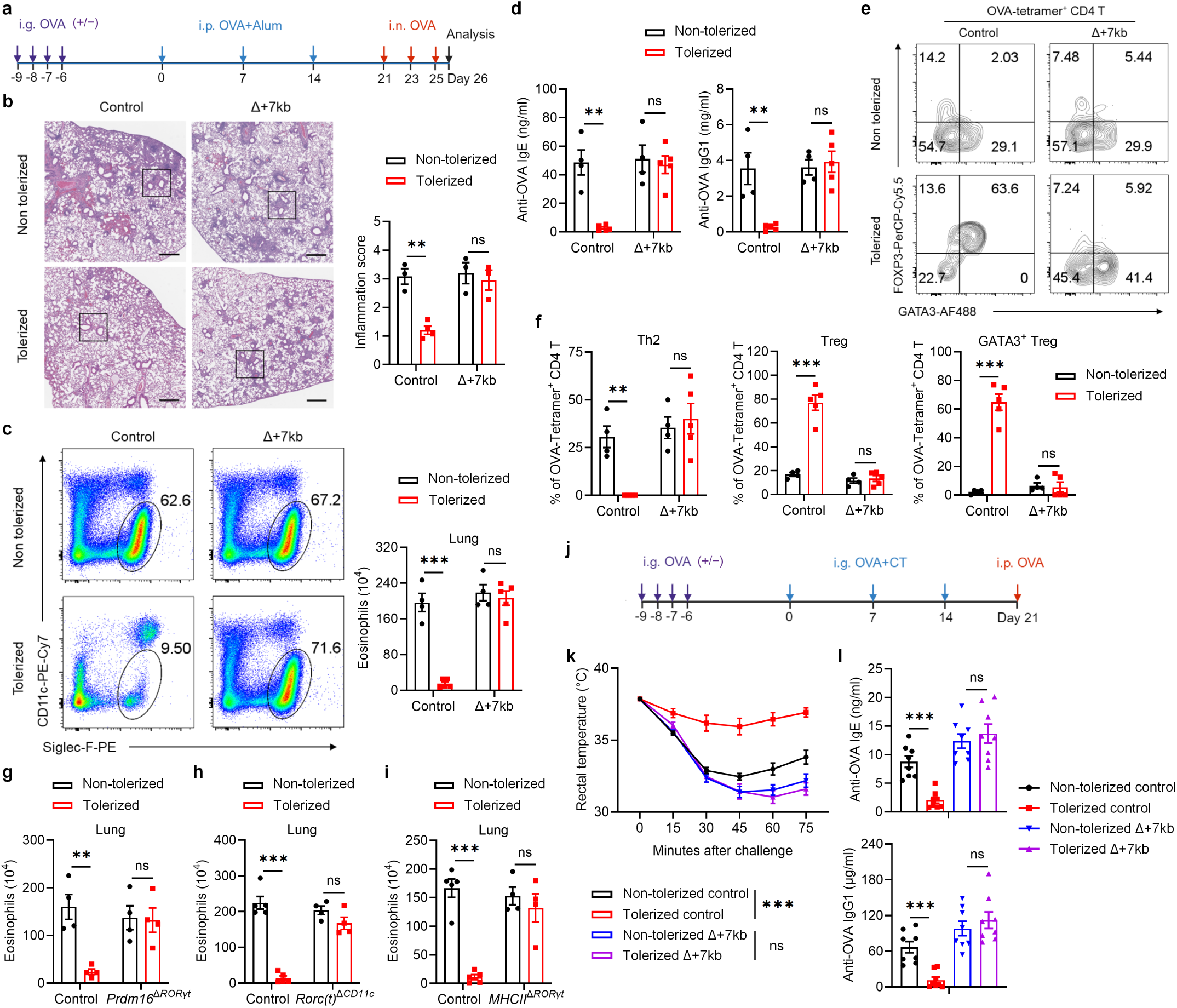
Tolerogenic DC are required to develop oral tolerance. **a**, Experimental design for the airway allergy experiments in **b**-**h**. i.g., intragastric; i.p., intraperitoneal; i.n., intranasal. **b**, Hematoxylin and eosin staining and inflammation score of the lung sections in control and Δ+7kb mice. Scale bars, 500 μm. Non-tolerized control mice, n = 3; tolerized control mice, n = 4; non-tolerized Δ+7kb mice, n = 3; tolerized Δ+7kb mice, n = 3. **c**-**f**, Eosinophil (CD45^+^CD11b^+^CD11c^low/−^Siglec-F^+^) numbers in the lung (**c**), OVA-specific IgE and IgG1 levels in the serum (**d**), phenotype of OVA:I-A^b^ tetramer^+^ CD4 T cells (**e**,**f**) in the lung of control and Δ+7kb mice. Flow cytometry plots in **e** were generated by concatenating the samples from each group. Non-tolerized control mice, n = 4; tolerized control mice, n = 5; non-tolerized Δ+7kb mice, n = 4; tolerized Δ+7kb mice, n = 5. **g**, Eosinophil numbers in the lung of control and *Prdm16*^Δ*RORγt*^ mice. n = 4. **h**, Eosinophil numbers in the lung of control and *Rorc(t)*^Δ*CD11c*^ mice. n = 4-5. **i**, Eosinophil numbers in the lung of control and *MHCII*^Δ*RORγt*^ mice. n = 4-5. **j**, Experimental design for the food allergy experiments in **k**,**l**. CT, cholera toxin. **k**,**l**, Changes in rectal temperature (**k**) and OVA-specific IgE and IgG1 levels in the serum (**l**) of control and Δ+7kb mice. n = 8. Data in **k**,**l** are pooled from two independent experiments. Data in **b-f** are representative of two independent experiments. Data are means ± s.e.m.; ns, not significant; statistics were calculated by unpaired two-sided t-test (**b**-**d**,**f**-**i**,**l**) and two-stage step-up method of Benjamini, Krieger, and Yekutieli (**k**).

To investigate whether tDC-induced pTregs are broadly required for tolerance to food antigens, we employed a food allergy model in which mice were either pre-fed or not fed OVA prior to sensitization intragastrically with OVA in cholera toxin (CT), followed by intraperitoneal OVA challenge (Fig. 4j). Systemic OVA challenge resulted in comparable anaphylactic responses, assessed as rapid reduction in core body temperature and elevated levels of serum OVA-specific IgE and IgG1, in tolerized Δ+7kb mice and non-tolerized Δ+7kb mice, whereas tolerized wild-type mice were protected (Fig. 4k,l). These findings indicate that tDC are crucial for the development of oral tolerance, highlighting their essential role in regulating immune responses to dietary antigens and preventing allergic responses.

## Orthologous tDC across human tissues

To determine whether the tDC identified in mice are also present in humans, we performed sc-RNA-seq on HLA-DR-enriched cells from freshly isolated mLNs of a 22-year-old trauma patient with no known history of atopy or chronic disease (Fig. 5a). Following the same unsupervised computation as for the murine experiments, we clustered the resulting single-cell dataset (Fig. 5b). A population of ILC3 was demarcated by expression of *IL7R*, *KIT*, and *NRP1* (Fig. 5c and Extended Data Fig. 10a). This was juxtaposed by an ILC1 cluster expressing *NCR1* and *TBX21*, as well as NK cells expressing *EOMES*, all consistent with prior literature^36^. A cluster positive for *XCR1* and *CLEC9A* was readily distinguished as cDC1. As compared to murine cDC2, there is less known about human cDC2 that allows for confident subset assignment. Nevertheless, we could identify a *SIRPA*^+^ population overlapping with *CD1C* expression, as would be expected for human cDC2^37^. We annotated a plasmacytoid DC cluster adjacent to this, as these cells were negative for *CD1C* while positive for *IL3RA* (CD123) and included rare cells positive for *TLR7* and *TLR9* expression^38^.

**Fig. 5.**
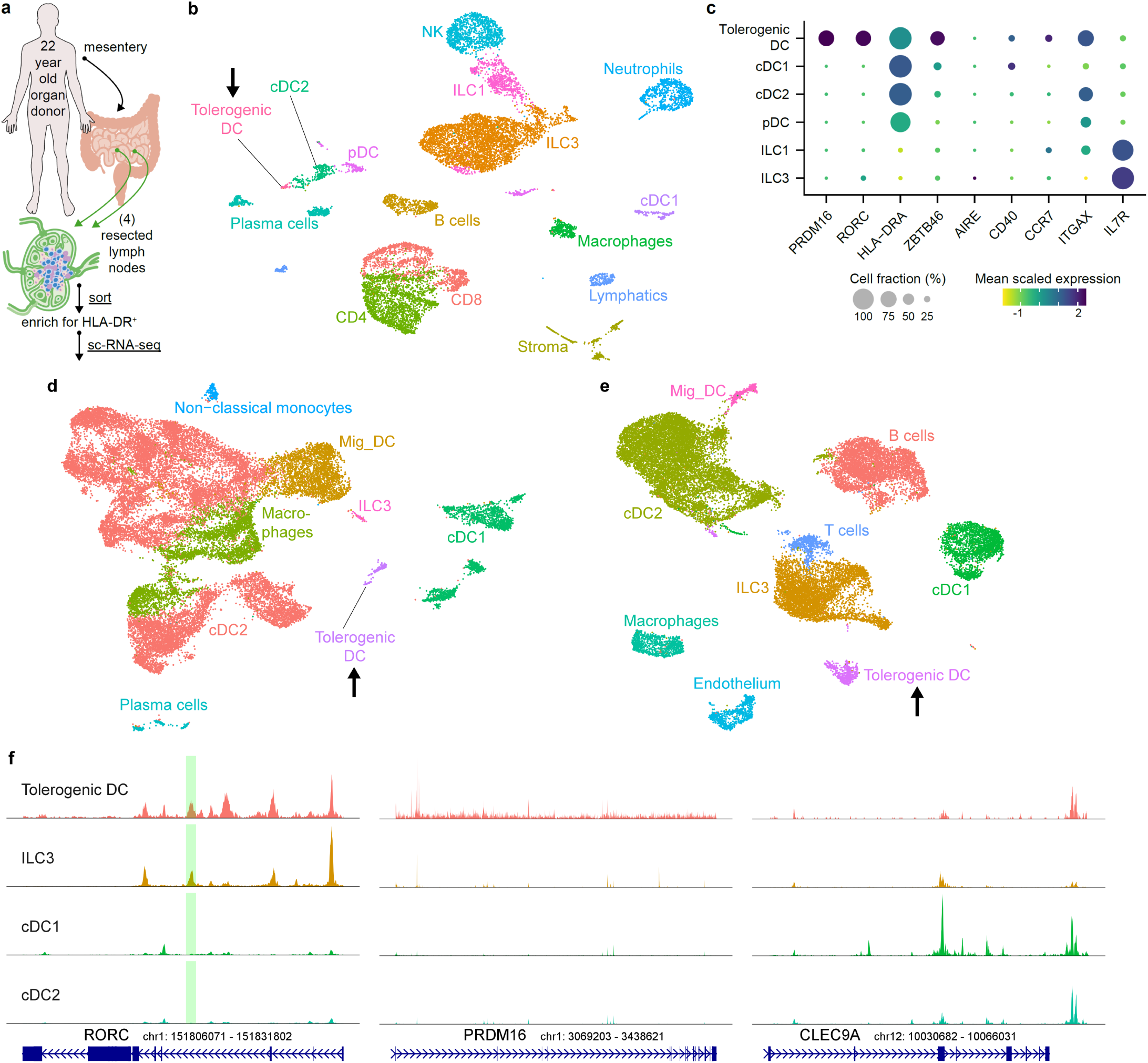
PRDM16-expressing tolerogenic DC are conserved in humans. **a**, Input scheme for human sc-RNA-seq, with four mLN resected and enriched for APCs. **b**, UMAP of 12,928 resultant human mLN transcriptomes. **c**, Dot plot of indicated clusters from **b** examining expression of genes previously ascribed to RORγt-APC subsets. **d**, UMAP derived from public datasets of human lamina propria (ileum and colon) resected from 6 donors, enriched for APCs. **e**, UMAP derived from public datasets of human tonsil from 9 donors, enriched for APCs. Black arrows in **b**,**d**,**e** highlight populations of tolerogenic DC. **f**, Chromatin accessibility profiles for *RORC*, *PRDM16*, and *CLEC9A* loci across the indicated cell types. Green shaded range demarcates human ortholog of murine *Rorc(t)* +7kb CRE, part of conserved non-coding sequence 9 (CNS9).

On querying for *RORC* in this human dataset, we observed non-ILC3 expression in a single set of cells clustering at the extremity of the cDC2 annotation (arrow, Fig. 5b,c). Strikingly, this same population exclusively displayed a high *PRDM16* signal. Unlike the murine sc-RNA-seq analysis, this population was not spontaneously demarcated by an unsupervised workflow, so we manually sub-clustered and similarly annotated it as tDC. This human cluster was also positive for *ZBTB46*, *CD40*, *CCR7*, and *ITGAX*, although *AIRE* was not detected in it or any other HLA-DR^+^ mLN population.

To further query for human tDC, we analyzed public datasets derived from other tissues. We assembled raw sequencing data from six colonic lamina propria and four paired ileal lamina propria digests (all resected >10cm from sites of colorectal cancer)^39^, as well as nine elective tonsillectomies^40,41^. Following our previous analysis, we annotated with special attention towards distinct populations with high co-expression of *PRDM16* and *RORC*. This revealed a clear tDC population in lamina propria (with equal contribution from ileum and colon, Fig. 5d), as well as tDC in tonsil (Fig. 5e). We examined multiple human bone marrow datasets but could not identify any cells co-expressing *PRDM16* and *RORC*. It therefore remains unclear when or where final tDC specification takes place.

We also examined available sc-ATAC-seq data, especially since murine *Rorc(t)* +7kb CRE has been demonstrated to have an ortholog contained within conserved non-coding sequence 9 of human *RORC*^20^. As in mouse, there was a prominent peak at this locus (Fig 5f, green shaded range) for human ILC3 and tDC. Similarly, *PRDM16* locus chromatin was accessible only in tDC, while *CLEC9A* was accessible in tDC and classical DC populations.

Aggregating all available data totaled 62,952 murine cells (including 1,271 tDC) and 67,638 human cells (including 988 tDC). By setting a relatively stringent definition of differentially expressed genes (DEGs) for each cell type within each dataset (Methods), we queried for DEGs shared across species. Conserved genes (Extended Data Fig. 10b, Supplemental Table 1) included mostly expected results for cDC1 (such as *CLEC9A*, *CADM1*, and *XCR1*) and ILC3 (such as *IL23R*, *IL22*, and *RORC*). This same analysis revealed 8 candidates conserved as differentially expressed genes across human and mouse tDC (Extended Data Fig. 10b,c), notably including *PRDM16*, *PIGR*, *IDO2*, *DLGAP1, MACC1*, and *CDH1*, largely corroborated by broader cross-species analyses of RORγt+ APCs^41,42^.

Genome-wide association studies have identified numerous single nucleotide polymorphisms (SNPs) of *PRDM16* having strong statistical association with autoimmune and inflammatory diseases, including asthma in African Americans (mostly pediatric)^43^, allergic rhinitis (IgE-mediated inflammation of the upper airway)^44^, rheumatoid arthritis^45^, and inflammatory bowel disease (IBD) in the Basque population^46^.

## Discussion

In maintaining intestinal homeostasis, the gut immune system is tasked with tolerating a complex array of dietary and microbial antigens, largely through the action of peripheral Tregs. Three independent studies demonstrated that RORγt-APCs, but not cDC, promote the differentiation of gut microbiota-specific pTregs. However, the precise identity of the pTreg-inducing RORγt-APCs remained unresolved. Here, we characterized the tolerogenic APCs as most closely resembling cDCs. Furthermore, our study showed that these APCs direct not only microbiota-, but also dietary antigen-specific pTreg cell differentiation, thus facilitating oral tolerance to food antigens. Oral tolerance can be enhanced by oral immunotherapy for food allergies^47,48^ and has been demonstrated effective in animal models to control antigen-specific autoimmune diseases^49,50^, raising the prospect that tDC could be harnessed to modulate autoimmune diseases and transplantation tolerance. Our findings indicate that the competency of tDC in establishing mucosal tolerance crucially relies on the transcription factors Prdm16 and RORγt, as well as a specific CRE within the *Rorc* locus. We previously showed that expression of MHCII in RORγt-expressing cells is sufficient for induction of pTreg cells^5^. Although the tDC are required for induction of pTreg cells, our studies don’t rule out the possibility that an additional RORγt^+^ APC also has a role in this differentiation process.

The transcriptional regulatory network in which Prdm16 and RORγt participate to influence development and function of these APCs remains to be elucidated. A comprehensive understanding of the key components will likely reveal human genetic variants that can predispose to inflammatory and allergic conditions. An understanding of the ontogeny of tDC may also provide critical insights into inflammatory diseases and potential therapeutic avenues. Crucially, these tolerogenic APCs possess a remarkable ability to induce pTregs despite their low abundance in the intestinal secondary lymphoid organs. It is intriguing that multiple genes conserved across human and mouse tDC are known for specialized cell adhesion and migration functions (*CDH1*, *MACC1*, *KRT8*, and *DLGAP1*). Unraveling the mechanisms behind their potent immunomodulatory effect is essential and will likely require elucidation of the spatial distribution of the APCs and temporal dynamics of differentiation of T cell subsets specific for antigens encountered in the alimentary tract. Notably, the abundance of these APCs is maintained from neonates to adulthood, consistent with previous^5,12,32^ and current findings that adult mice retain the capacity to induce tolerance to microbiota and oral antigen. The tDC may therefore serve as a valuable therapeutic target to enhance Treg functions in multiple immune-related diseases.

## Acknowledgements

We thank members of the Littman laboratory for valuable discussion and critical reading of the manuscript; S. R. Schwab and M. Okuniewska for providing OT-II;UBC-GFP;*Rag1*^−/−^ mice; B. Spiegelman for providing *Prdm16*^fl/fl^ mice; M. Oukka for providing *Il23r*^gfp^ mice. S. Y. Kim and the NYU Rodent Genetic Engineering Laboratory for assistance with generation and rederivation of mutant mice; the Genome Technology Core for scRNA sequencing (RRID: SCR_017929); C. Loomis and the Experimental Pathology Research Laboratory for histology of tissue samples; and the NIH Tetramer Core Facility for generating MHC class II tetramers. NYU core facilities are partially supported by NYU Cancer Center Support Grant P30CA016087 at the Laura and Isaac Perlmutter Cancer Center. L.F. is supported by a Cancer Research Institute Irvington Postdoctoral Fellowship (CRI12690). R.U. is supported by a Clinical Scientist Career Development Award (K08CA283272), Stand Up to Cancer Phillip A. Sharp Innovation in Collaboration Award (SU2C-AACR-PS-36), and by the Rosenfield & Glassman Foundation. F.M.C. is supported by the National Institutes of Health (T32AL100853, F32AI181496). This work was supported by R01AI158687 (D.R.L.), and the Howard Hughes Medical Institute (D.R.L.).

## Author contributions

L.F., R.U. and D.R.L. conceived the project. L.F designed and performed most mouse experiments and analyzed the data. M.P. and F.M.C. performed mouse experiments and analyzed the data. G.R.M. assisted with mouse experiments. M.P. performed bulk ATAC-sequencing and analyzed the data. M.P. generated *Rorc(t)* cis-regulatory element mutant mice. L.F., and R.U. and G.R.M. analyzed histology. L.F. performed sc-RNA seq and multiome scRNA-seq/scATAC-seq of mouse samples and contributed to computational analysis. R.U. performed sc-RNA seq of human samples. A.G. arranged and acquired human tissues. R.U. performed all scRNA-seq and scATAC-seq computational analysis. L.F., R.U. and D.R.L. wrote the manuscript, with input from the other authors. D.R.L. supervised the research and contributed to experimental design.

## Data and code availability

Mouse and human RNA-seq and ATAC-seq data generated for this project, along with all code used for computational analysis, will be made available for open-access download at: https://zenodo.org/record/xxx.

---

**Extended Data Fig. 1.**
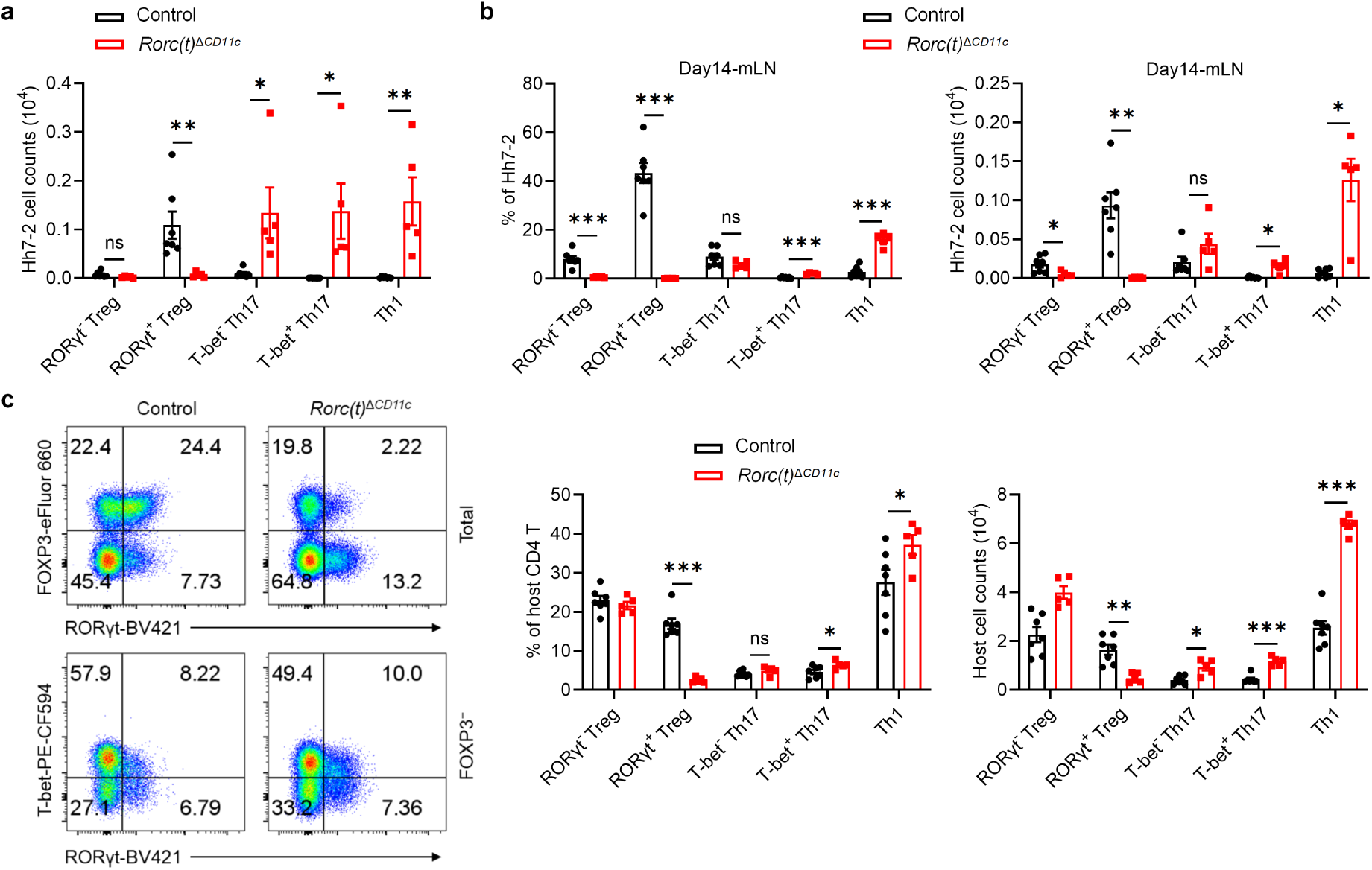
RORγt expression in CD11c lineage APC is necessary for directing the differentiation of gut microbiota-specific pTregs. **a**, Numbers of *Hh*-specific pTreg, Th17 and Th1 cells in the LILP of *Hh*-colonized control (n = 7) and *Rorc(t)*^Δ*CD11c*^ (n = 5) mice at 14 days after adoptive transfer of naïve Hh7-2tg CD4^+^ T cells. **b**, Phenotype of *Hh*-specific T cells in the mLN of mice shown in **a**. **c**, Phenotype of host CD4 T cells in the LILP of mice shown in **a**, with representative flow cytometry profiles (left) and aggregate quantitative data (right). The flow cytometry plots are gated on total (upper) and FOXP3^−^ (lower) host CD4 T cells (CD45^+^B220^−^TCRγδ^−^TCRβ^+^CD4^+^CD90.1^−^). Data are pooled from two independent experiments. Data are means ± s.e.m.; ns, not significant; statistics were calculated by unpaired two-sided t-test.

**Extended Data Fig. 2.**
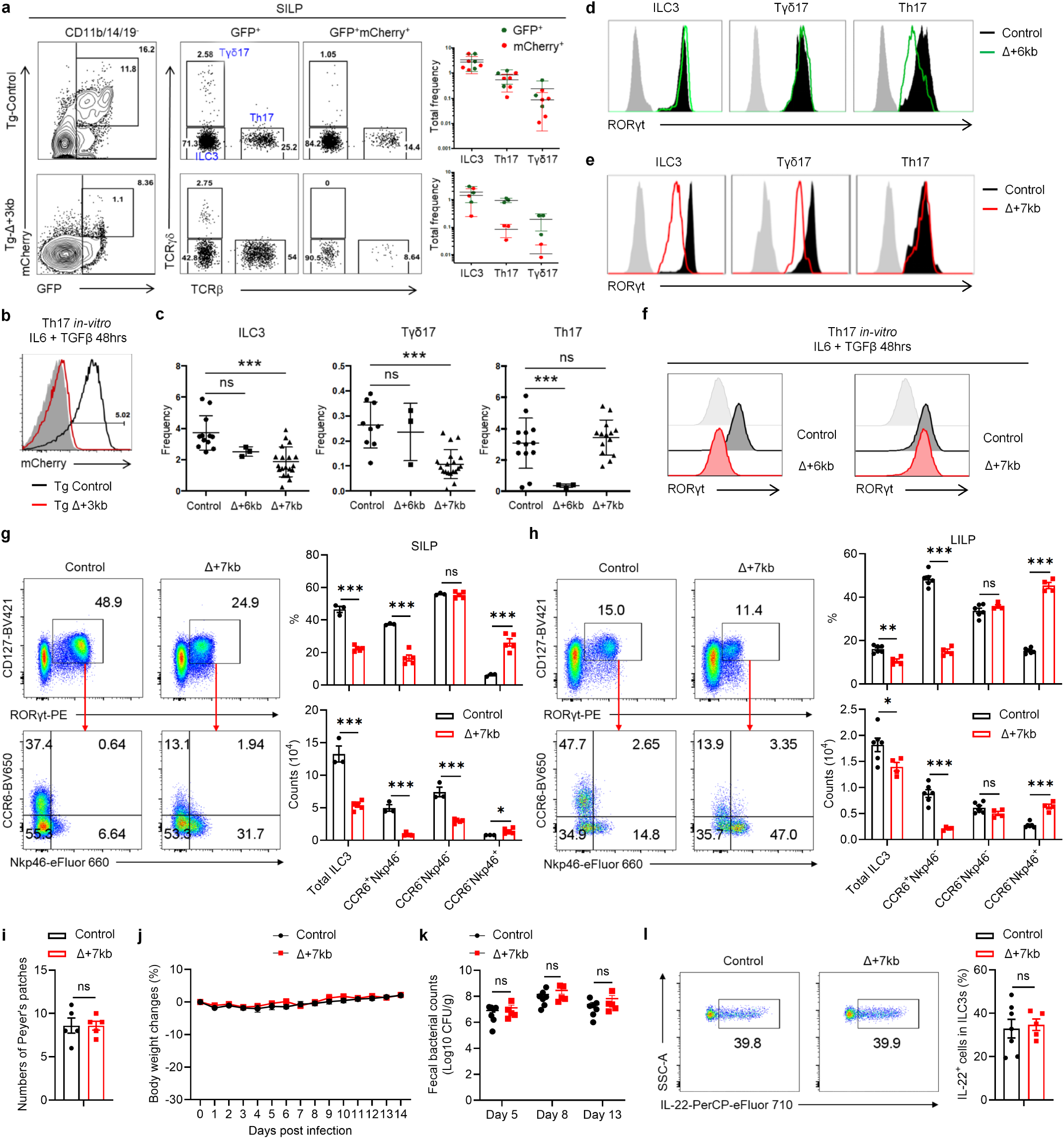
Characterization of lineage-specific *Rorc(t)* cis-regulatory elements. **a**,**b**, Representative flow cytometry plots and summary graphs depicting comparison of SILP GFP^+^ and mCherry^+^ populations (**a**) as well as mCherry expression of *in vitro* differentiated Th17 cells (**b**) in BAC transgenic mice bred to RORγt GFP reporter mice (heterozygous for gfp knockout allele), Tg (Control *Rorc(t)*-mCherry);*Rorc(t)*^+/gfp^ and Tg (Δ+3kb *Rorc(t)*-mCherry);*Rorc(t)*^+/gfp^ mice. **c**, Summary of indicated SILP RORγt^+^ populations as frequency of SILP mononuclear cells from control, Δ+6kb and Δ+7kb mice. **d**,**e**, Representative histograms showing RORγt expression in the SILP RORγt^+^ populations from mice shown in **c**. **f**, RORγt expression among *in vitro* differentiated Th17 cells isolated from mice shown in **c**. **g**,**h**, Phenotype of ILC3 (CD45^+^Lin^−^CD127^+^RORγt^+^) subsets in the SILP (**g**) and LILP (**h**) of control and Δ+7kb mice. SILP: control mice, n = 3; Δ+7kb mice, n = 5. LILP: control mice, n = 6; Δ+7kb mice, n = 4. **i**, Numbers of Peyer’s patches in control (n = 5) and Δ+7kb (n = 5) mice. **j**-**l**, Body weight changes (**j**), fecal *C. rodentium* counts (**k**) and frequencies of IL-22^+^ cells in RORγt^+^ ILC3 (day 14, *ex vivo* stimulation with IL-23) in the LILP (**l**) of control (n = 7) and Δ+7kb (n = 5) mice post *C. rodentium* infection. Data in **j**-**l** are pooled from two independent experiments. Data in **g**-**i** are representative of two (**g**,**h**) or three (**i**) independent experiments. Data are means ± s.d.; ns, not significant; statistics were calculated by unpaired two-sided t-test.

**Extended Data Fig. 3.**
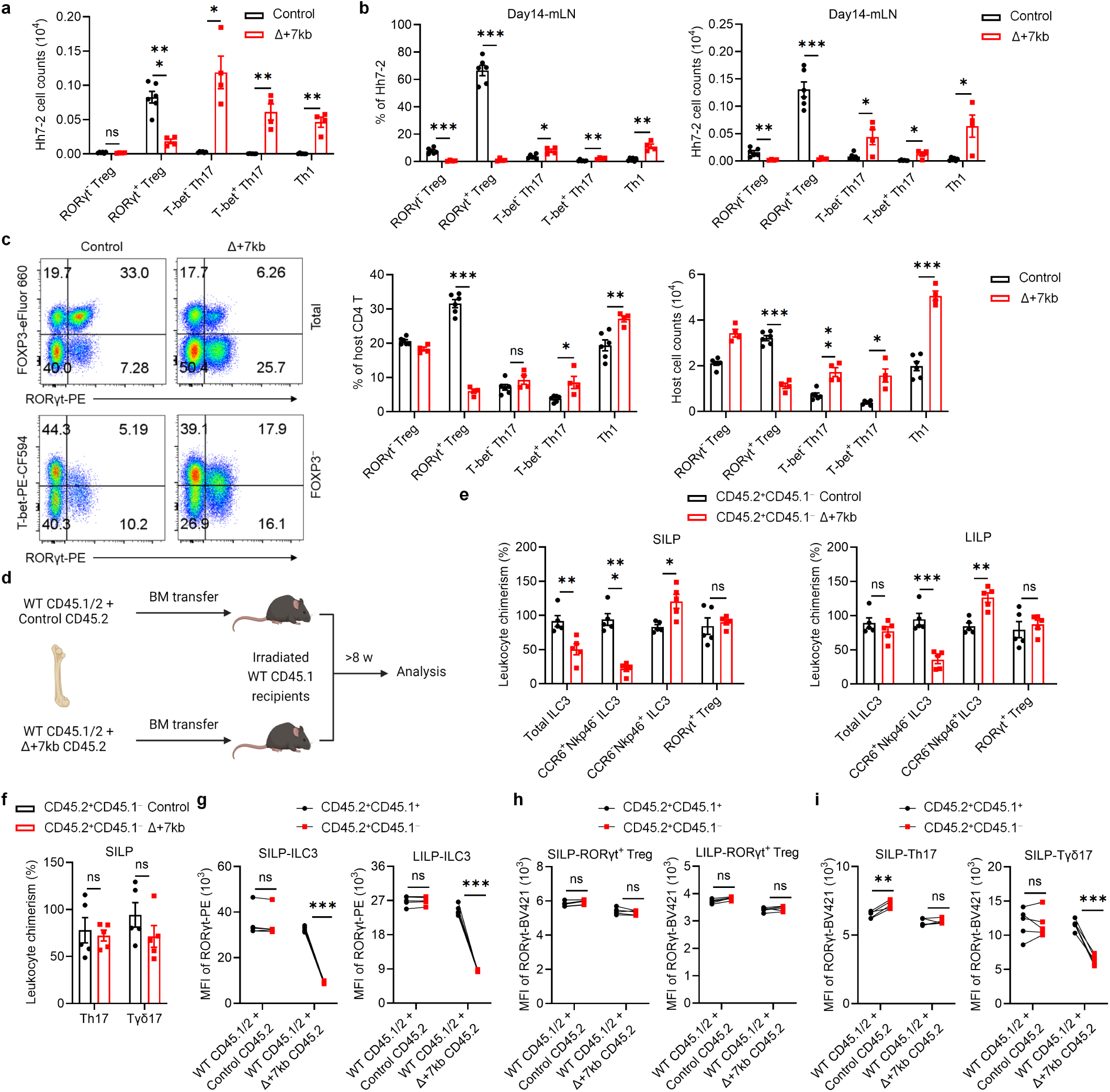
*Rorc(t)* +7kb regulates RORγt^+^ Treg in a cell-extrinsic manner. **a**, Numbers of *Hh*-specific T cells in the LILP of *Hh*-colonized control (n = 6) and Δ+7kb (n = 4) mice at 14 days after adoptive transfer of naïve Hh7-2tg CD4^+^ T cells. **b**, Phenotype of *Hh*-specific T cells in the mLN of mice shown in **a**. **c**, Phenotype of host CD4 T cells in the LILP of mice shown in **a**. **d**, Experimental design for the bone marrow (BM) chimeric experiments in **e**-**i**. **e**,**f**, Relative CD45.2^+^CD45.1^-^ leukocyte chimerism normalized to CD45.2^+^CD45.1^−^ splenic B cells (n = 5). **g**-**i**, RORγt mean fluorescence intensity (MFI) of CD45.2^+^CD45.1^+^ and CD45.2^+^CD45.1^−^ RORγt^+^ cells in the SILP and LILP (n = 5). Data are representative of two independent experiments. Data are means ± s.e.m.; ns, not significant; statistics were calculated by unpaired two-sided t-test (**a**-**c**,**e**,**f**) and paired two-sided t-test (**g**-**i**).

**Extended Data Fig. 4.**
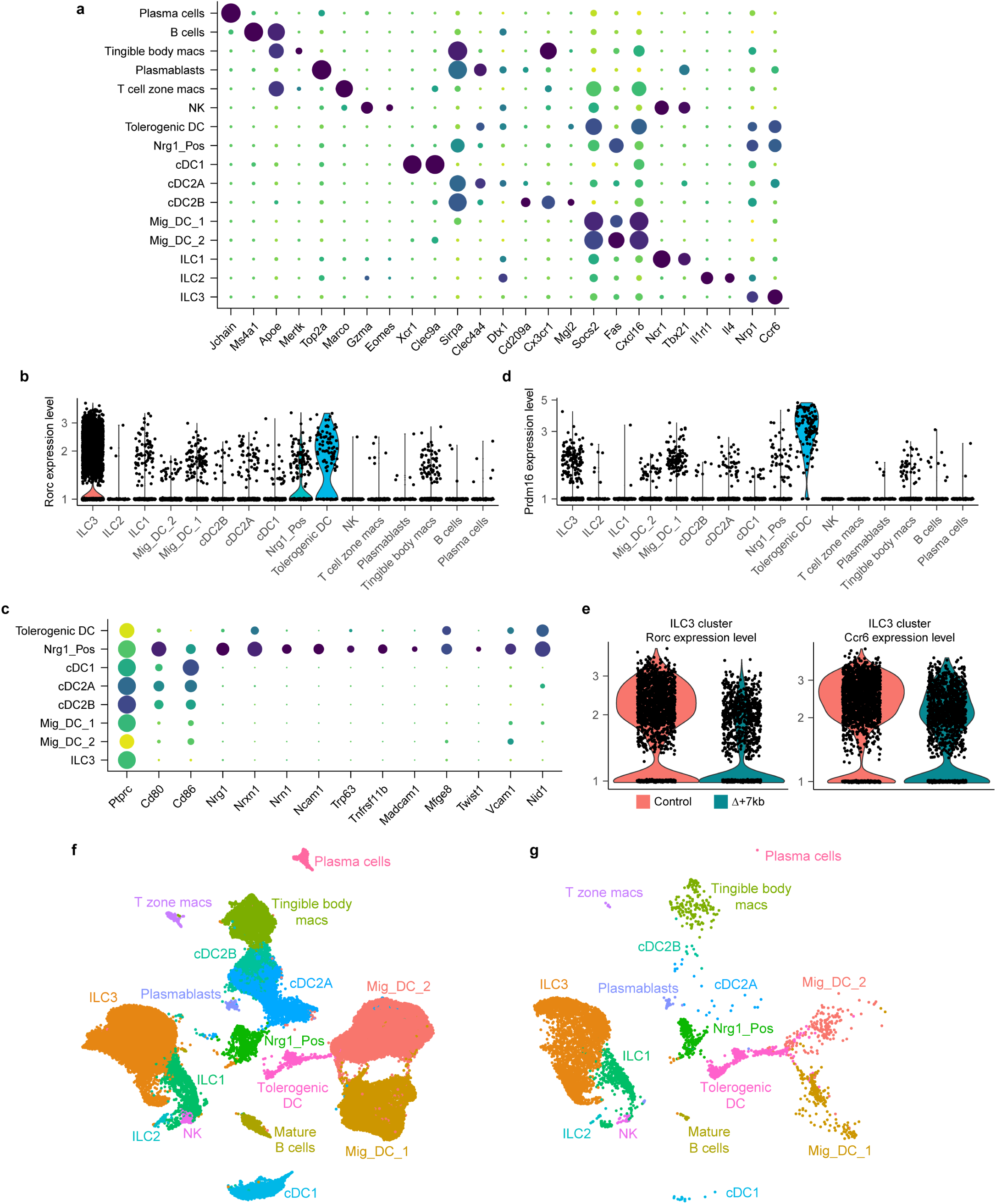
Sequencing annotation and analysis of Δ+7kb mouse model mLN and multiome datasets. **a**, Dot plot of all 16 cell types from Δ+7kb mouse model mLN (mutant and control mice combined), demonstrating canonical genes used to annotate each cluster. **b**, Violin plot of *Rorc* expression across all clusters. **c**, Dot plot of select APC clusters, examining genes described for Nrg1_Pos as well as FRC/mTEC cell types. **d**, Violin plot of *Prdm16* expression across all clusters. **e**, Violin plots of *Rorc* and *Ccr6* expression within the ILC3 cluster, comparing Δ+7kb mutant versus control biological conditions. **f**, Annotated UMAP with datasets combined from all murine experiments (all mutant and control animals, as well subsequent multi-ome experiment). **g**, Re-analysis of raw data from Akagbosu *et al*., which was computationally integrated alongside data in **f**.

**Extended Data Fig. 5.**
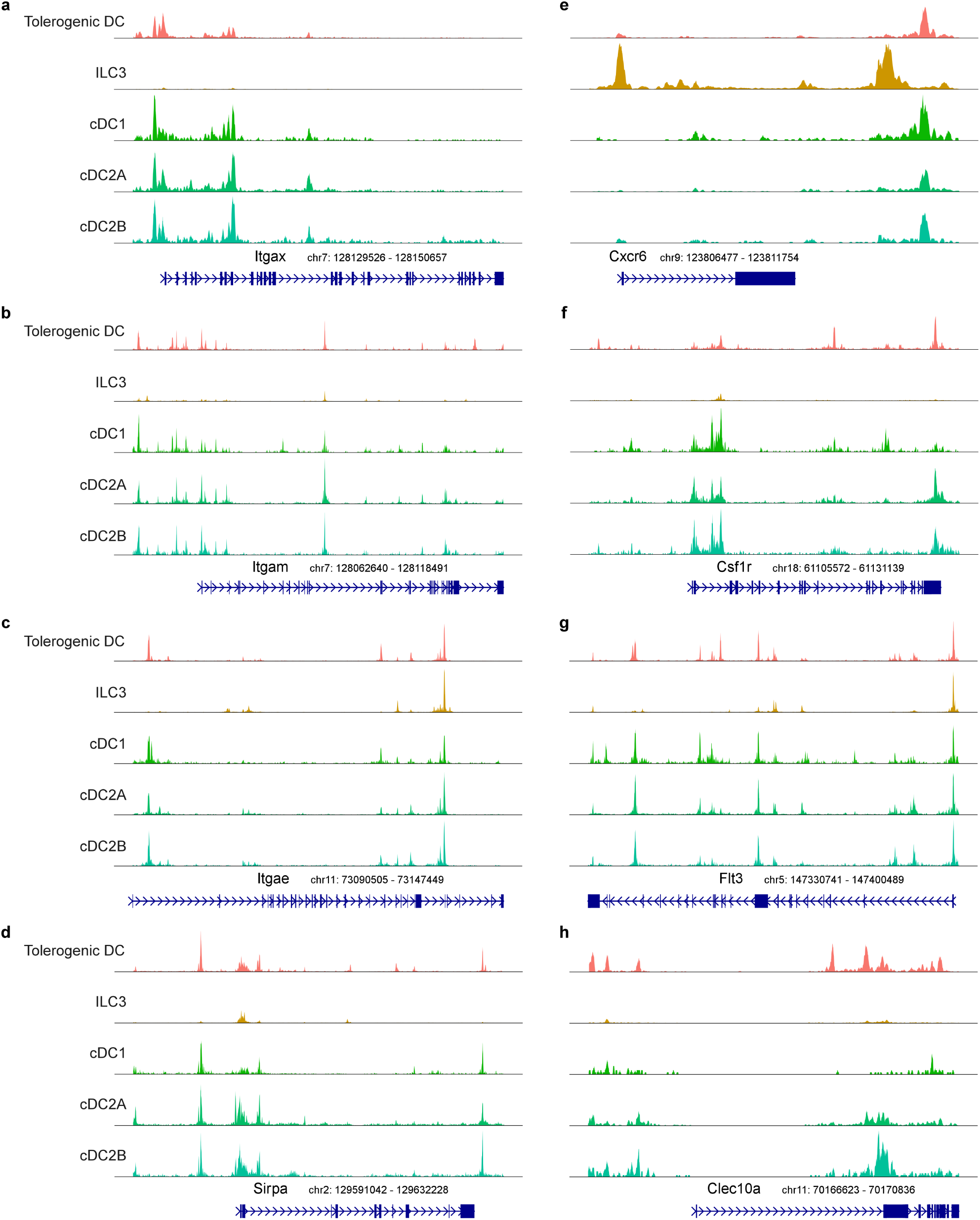
Comparing the epigenetic landscape across APC populations. **a-h,** Chromatin accessibility profiles for *Itgax*, *Itgam*, *Itgae*, *Sirpa*, *Cxcr6*, *Csf1r*, *Flt3*, and *Clec10a* loci across the indicated mouse APC populations.

**Extended Data Fig. 6.**
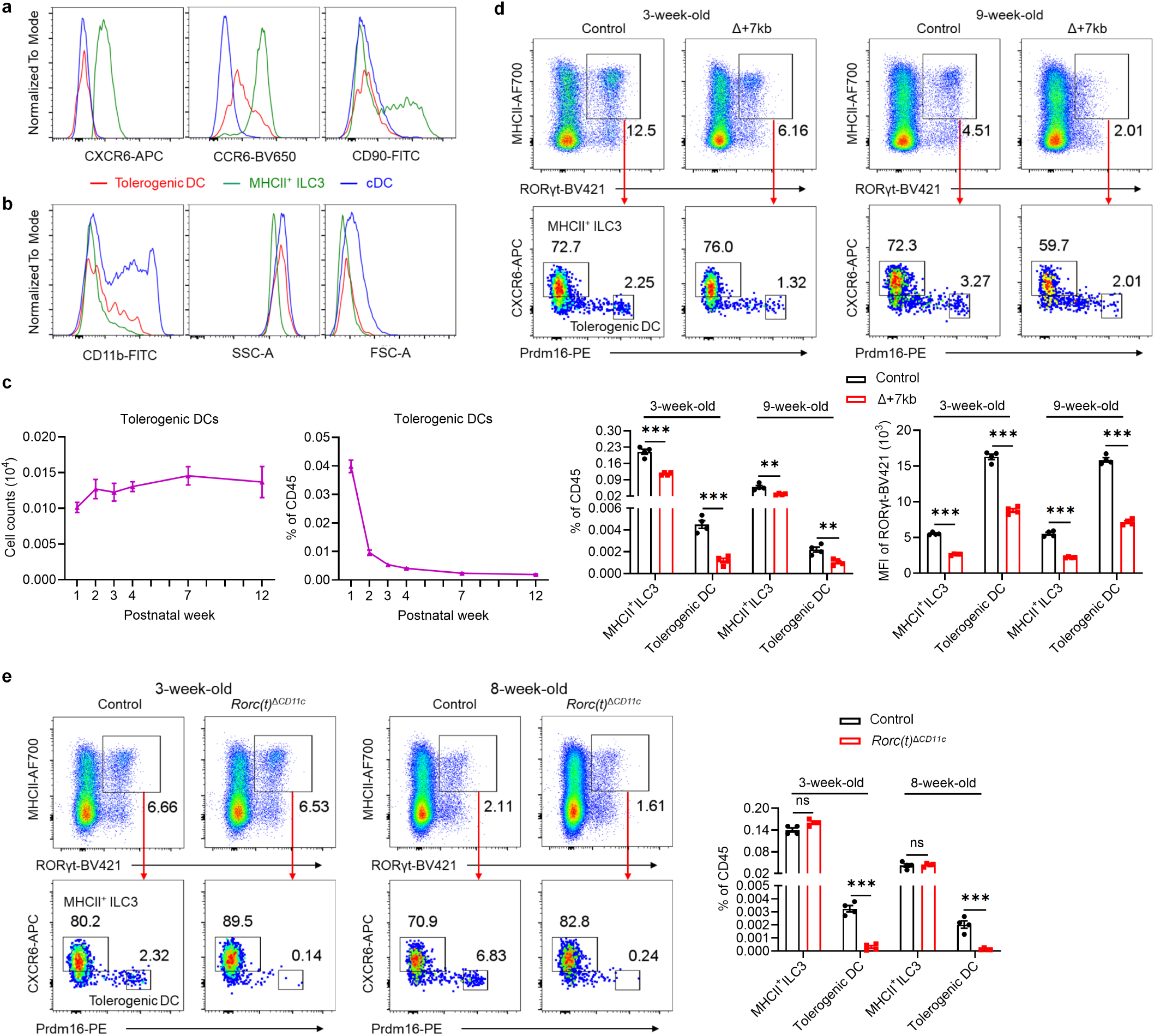
Selective loss of tolerogenic DC in *Rorc(t)*^Δ*CD11c*^ mice. **a**,**b**, Expression of the indicated proteins in tolerogenic DC, MHCII^+^ ILC3 and cDC, as gated in Fig. 2f. **c**, Numbers and frequencies of tolerogenic DC (CD45^+^Ly6G^−^B220^−^TCRγδ^−^TCRβ^−^MHCII^+^RORγt^+^CXCR6^−^Prdm16^high^) in mLN from week 1 to week 12 (n = 5-6 per timepoint). **d**, Representative flow cytometry plots (top), frequencies (bottom left) and RORγt MFI (bottom right) of tolerogenic DC and MHCII^+^ ILC3 in the mLN of 3-week-old and 9-week-old control (n = 4) and Δ+7kb (n = 4) mice. **e**, Representative flow cytometry plots (left) and frequencies (right) of tolerogenic DCs and MHCII^+^ ILC3 in the mLN of 3-week-old and 8-week-old control (n = 4) and *Rorc(t)*^Δ*CD11c*^ (n = 4) mice. The top flow cytometry plots are gated on CD45^+^Ly6G^−^B220^−^TCRγδ^−^TCRβ^−^. Data in **a**,**b**,**d**,**e** are representative of two (**d**,**e**) or three (**a**,**b**) independent experiments. Data are means ± s.e.m.; ns, not significant; statistics were calculated by unpaired two-sided t-test.

**Extended Data Fig. 7.**
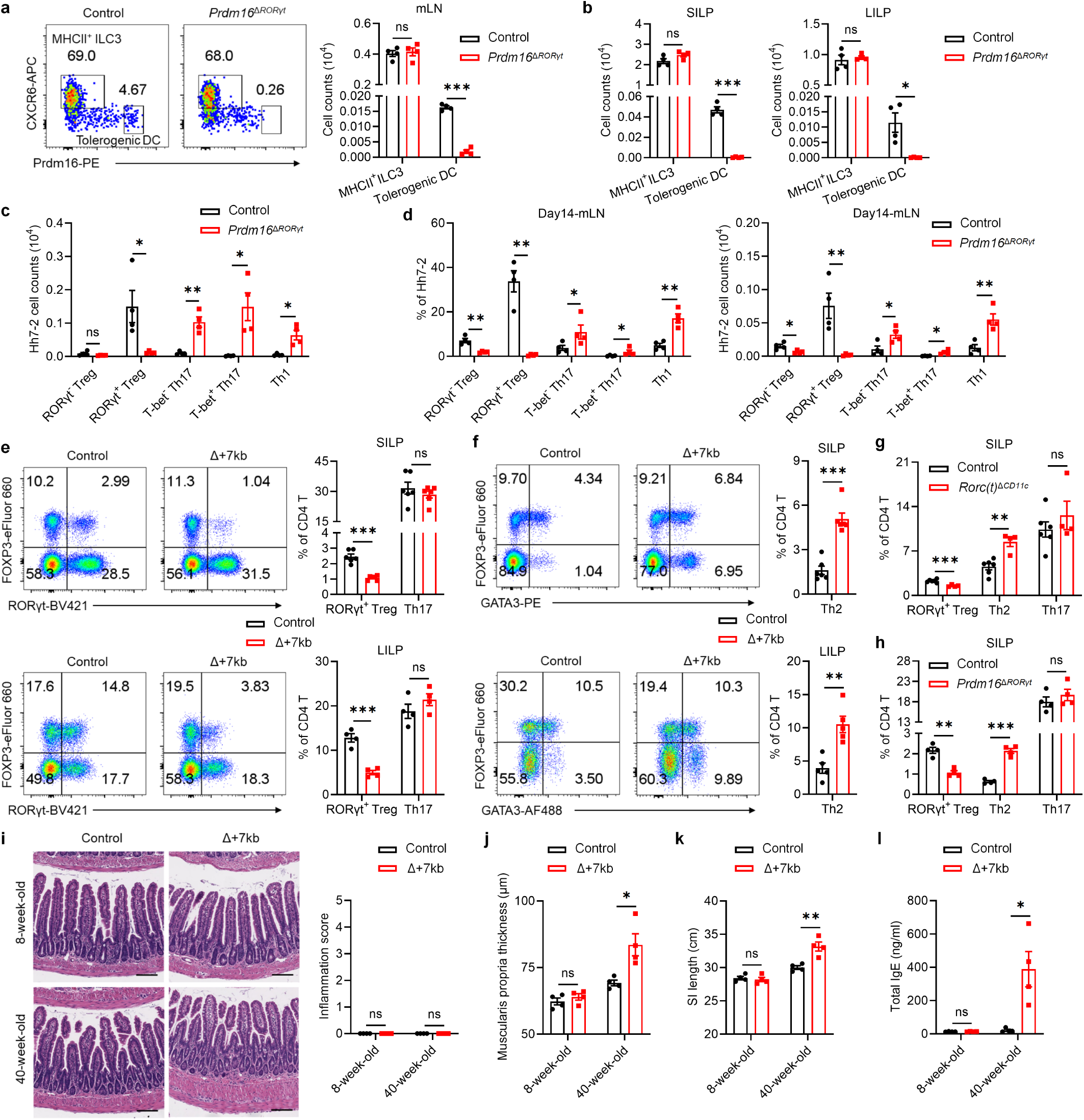
Tolerogenic DC deficiency leads to type 2 gastrointestinal pathology. **a**,**b**, Representative flow cytometry plots and numbers of tolerogenic DC and MHCII^+^ ILC3 in the mLN (**a**), SILP and LILP (**b**) of control (n = 4) and *Prdm16*^Δ*RORγt*^ (n = 4) mice. The flow cytometry plots are gated on CD45^+^Ly6G^−^B220^−^TCRγδ^−^TCRβ^−^MHCII^+^RORγt^+^. **c**, Numbers of *Hh*-specific T cells in the LILP of *Hh*-colonized control (n = 4) and *Prdm16*^Δ*RORγt*^ (n = 4) mice at 14 days after adoptive transfer of naïve Hh7-2tg CD4^+^ T cells. **d**, Phenotype of *Hh*-specific T cells in the mLN of mice shown in **c**. **e**,**f**, Representative flow cytometry plots and frequencies of RORγt^+^ Treg, Th17 and Th2 cells in the SILP and LILP of control (n = 4-6) and Δ+7kb mice (n = 4-6). **g**,**h**, Frequencies of RORγt^+^ Treg, Th2 and Th17 cells in the SILP of control and mutant mice. **g**, control mice, n = 6; *Rorc(t)*^Δ*CD11c*^ mice, n = 4. **h**, control mice, n = 4; *Prdm16*^Δ*RORγt*^ mice, n = 4. **i**,**j**, Hematoxylin and eosin staining and inflammation score (**i**), and average muscularis propria thickness (**j**) of the distal small intestine sections in 8-week-old and 40-week-old control (n = 4) and Δ+7kb (n = 4) mice. Scale bars, 100 μm. **k**,**l**, Small intestine length (**k**) and serum total IgE levels (**l**) of 8-week-old and 40-week-old control (n = 4) and Δ+7kb (n = 4) mice. Data are representative of two (**a**-**d**,**g**,**h**) or three (**e**,**f**) independent experiments. Data are means ± s.e.m.; ns, not significant; statistics were calculated by unpaired two-sided t-test.

**Extended Data Fig. 8.**
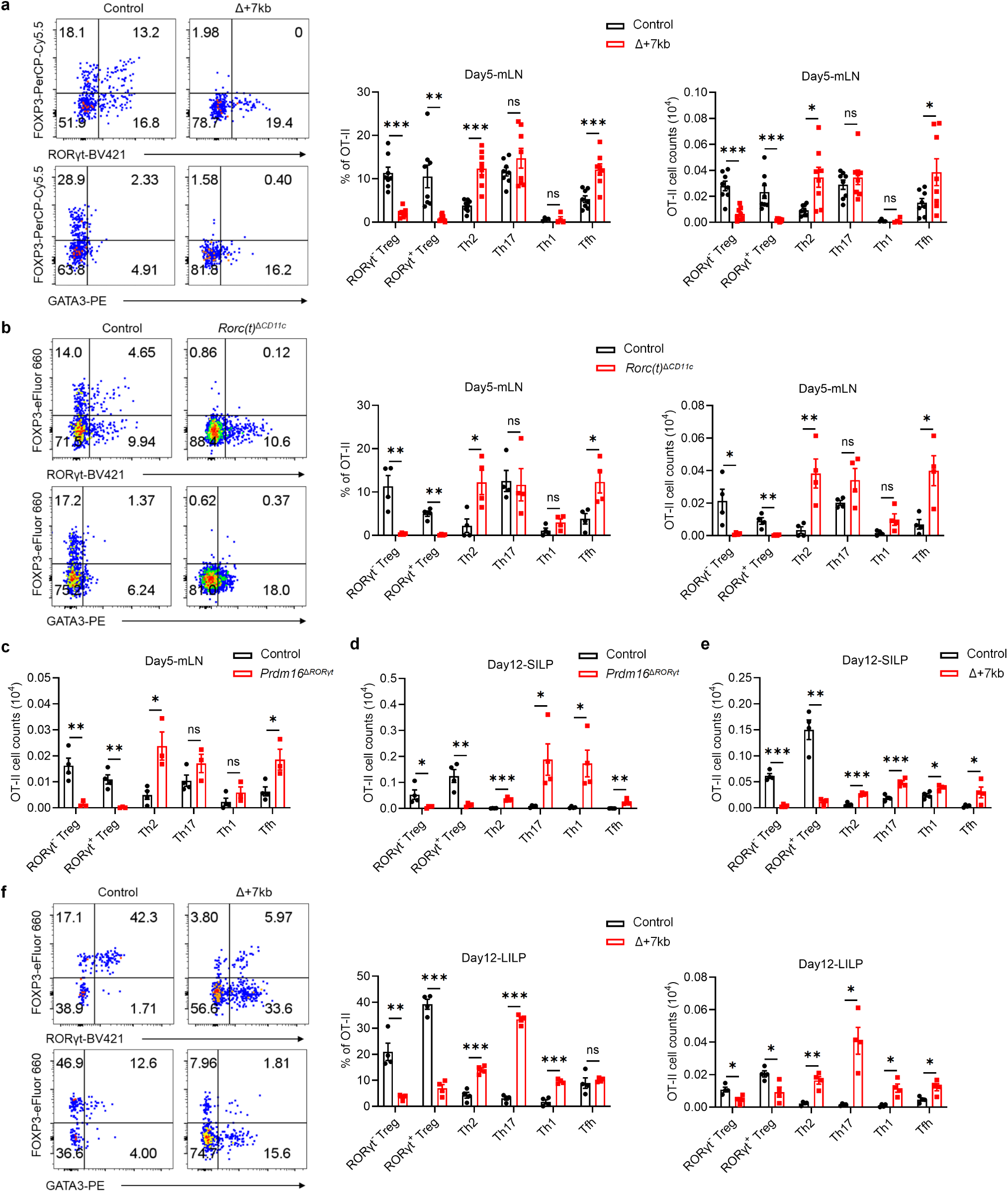
Tolerogenic DC are essential for the differentiation of food antigen-specific pTregs. **a-e**, Phenotype of OT-II pTreg, Th2, Th17, Th1 and Tfh cells in the mLN, SILP and LILP of OVA-fed control and mutant mice at 5 and 12 days post-adoptive transfer of naïve OT-II CD4^+^ T cells. **a**, mLN: control mice, n = 8; Δ+7kb mice, n = 8. **b**, mLN: control mice, n = 4; *Rorc(t)*^Δ*CD11c*^ mice, n = 4. **c**, mLN: control mice, n = 4; *Prdm16*^Δ*RORγt*^ mice, n = 3. **d**, SILP: control mice, n = 4; *Prdm16*^Δ*RORγt*^ mice, n = 4. **e**, SILP: control mice, n = 4; Δ+7kb mice, n = 4. **f**, LILP: control mice, n = 4; Δ+7kb mice, n = 4. Data in **a** are pooled from two independent experiments. Data in **b**,**d-f** are representative of two (**b**,**d**) or three (**e**,**f**) independent experiments. Data are means ± s.e.m.; ns, not significant; statistics were calculated by unpaired two-sided t-test.

**Extended Data Fig. 9.**
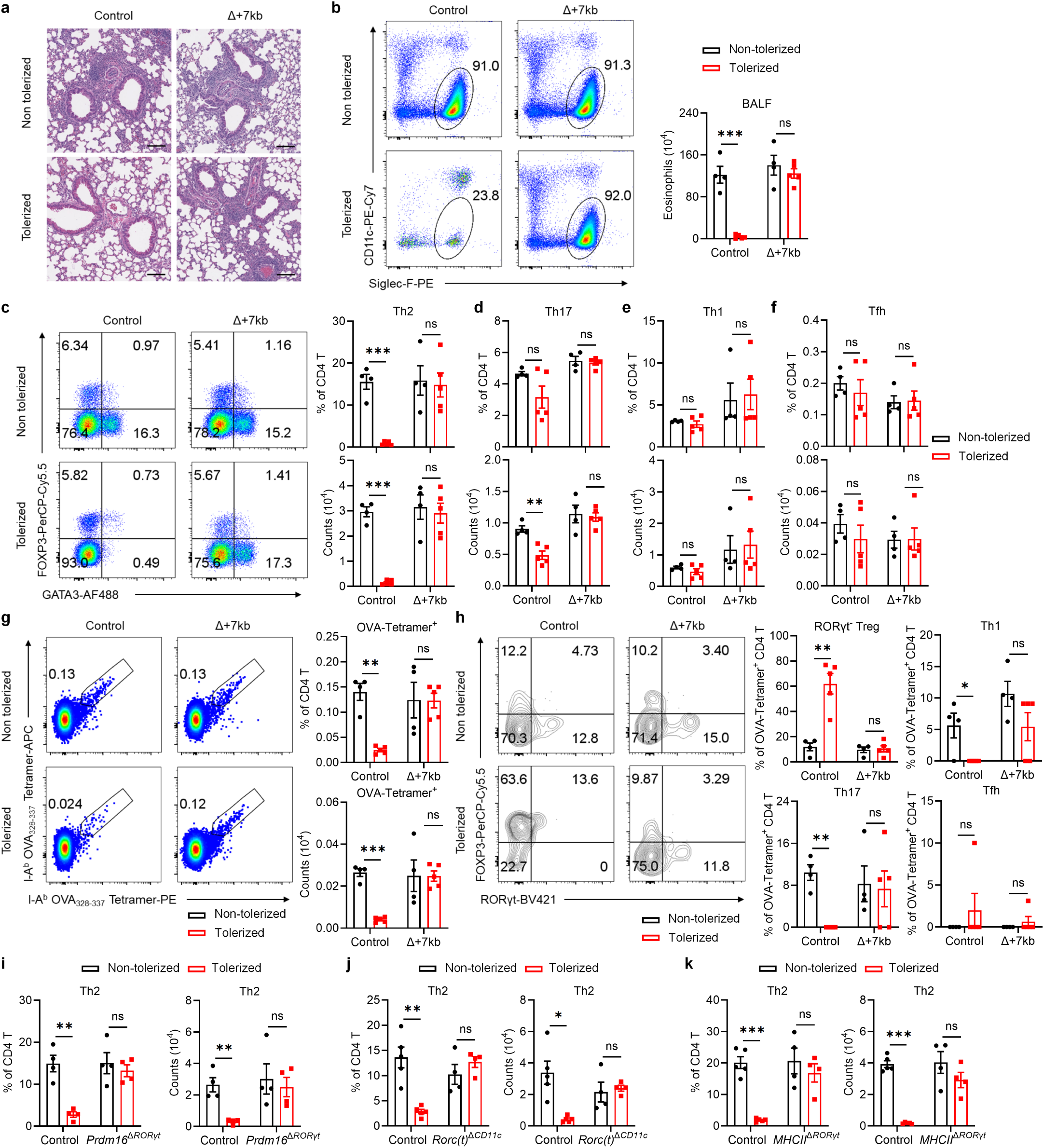
Tolerogenic DC are required for establishing oral tolerance against allergic airway responses. **a**, Magnified images of the lung sections in Fig. 4b. Scale bars, 100 μm. **b**-**h**, Eosinophil numbers in the BALF (bronchoalveolar lavage fluid) (**b**), phenotype of total CD4 T cells (**c**-**f**) and OVA:I-A^b^ tetramer^+^ CD4 T cells (**g,h**) in the lung of control and Δ+7kb mice shown in Fig. 4c-**f**. Flow cytometry plots in **g,h** were generated by concatenating the samples from each group. **i**, Phenotype of total Th2 cells in the lung of control and *Prdm16*^Δ*RORγt*^ shown in Fig. 4g. **j**, Phenotype of total Th2 cells in the lung of control and *Rorc(t)*^Δ*CD11c*^ shown in Fig. 4h. **k**, Phenotype of total Th2 cells in the lung of control and *MHCII*^Δ*RORγt*^ shown in Fig. 4i. Data in **b-h** are representative of two independent experiments. Data are means ± s.e.m.; ns, not significant; statistics were calculated by unpaired two-sided t-test.

**Extended Data Fig. 10.**
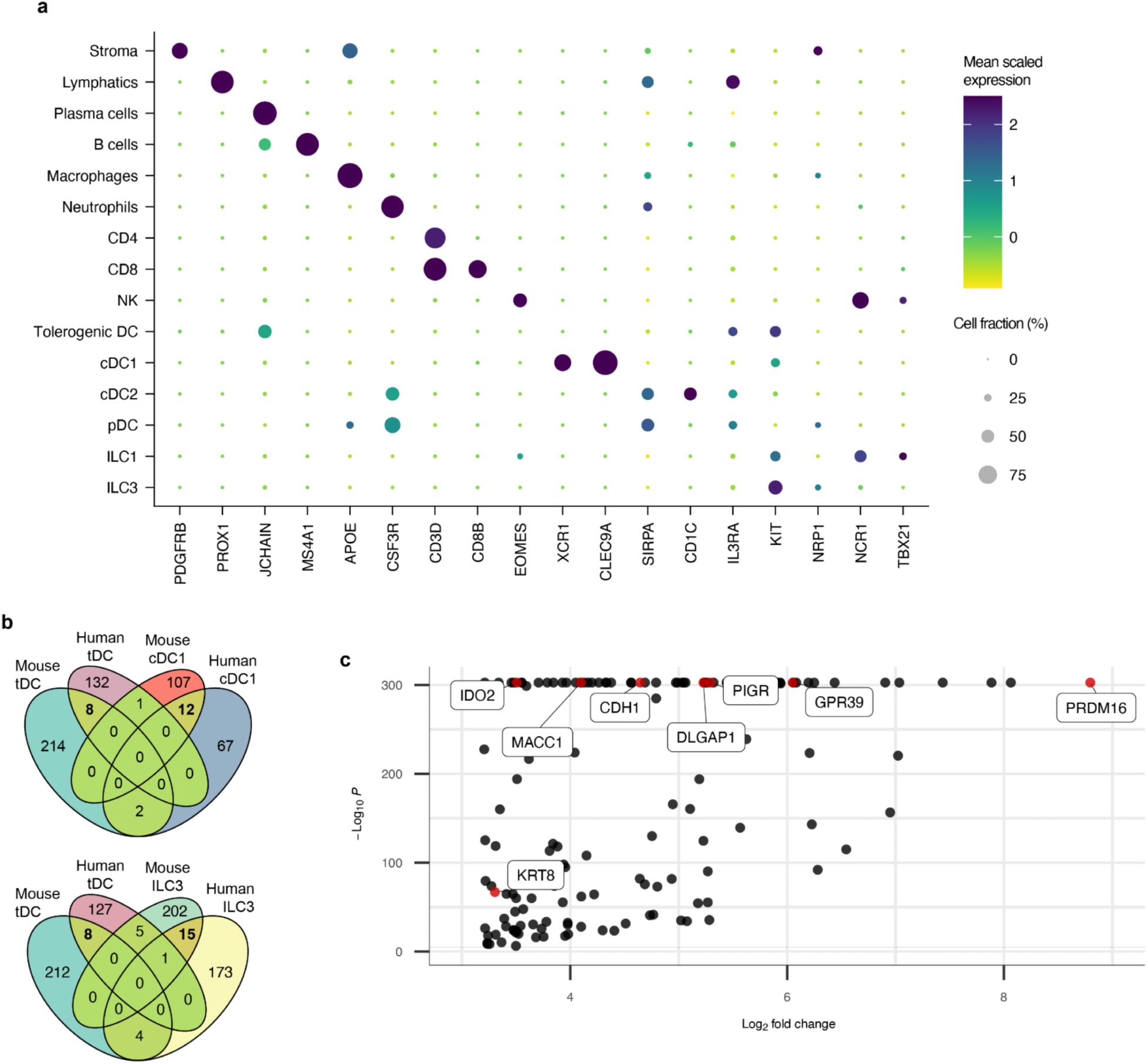
Analysis of differentially upregulated genes shared by human and mouse tDC. **a**, Dot plot of all 15 cell types from human mLN sc-RNA-seq experiment, demonstrating canonical genes used to annotate each cluster. **b**, Enumeration of differentially upregulated genes above a threshold of Log_2_(Fold Change) = 3.2 after integrating all available data, from all tissues, for indicated murine and human APC populations. Overlaps within the Venn diagrams demonstrate conserved genes. **c**, Volcano plot of all 141 genes differentially upregulated in human tDC, with explicit annotation of the 8 genes shared with mouse tDC, as seen in **b**.

## Methods

### Mice

C57BL/6 mice (Jax 000664), CD45.1 mice (B6.SJL-*Ptprc^a^ Pepc^b^*/BoyJ, Jax 002014), CD90.1 mice (B6.PL-*Thy1^a^*/CyJ, Jax 000406), *Cd11c*^cre^ mice (B6.Cg-Tg(Itgax-cre)1-1Reiz/J, Jax 008068), *I-AB*^fl/fl^ mice (B6.129X1-*H2-Ab1^b-tm1Koni^*/J, Jax 013181) and *ROSA26*^lsl-tdTomato/lsl-tdTomato^ mice (B6.Cg-*Gt(ROSA)26Sor^tm14(CAG-tdTomato)Hze^*/J, Jax 007914) were purchased from the Jackson Laboratories. *Rorc*^fl/fl^, *Rorc(t)*^gfp/gfp^, *Rorc(t)*^cre^, and Hh7-2tg mice were generated in our laboratory and have been described^15,32,51,52^. *Il23r*^gfp^ mice^53^ were provided by M. Oukka. OT-II;UBC-GFP;*Rag1*^−/−^ mice^54–56^ were provided by S. R. Schwab. *Prdm16*^fl/fl^ mice^57^ were provided by B. Spiegelman. Tg (Δ+3kb *Rorc(t)*-mCherry), *Rorc(t)* +6kb^−/−^and *Rorc(t)* +7kb^−/−^ mice were generated as described in ‘Generation of BAC transgenic reporter and CRISPR knockout mice’. Mice were bred and maintained in the Alexandria Center for the Life Sciences animal facility of the New York University School of Medicine, in specific pathogen-free conditions. Sex- and age-matched mice used in all experiments were 6-12 weeks old at the starting point of treatment if not otherwise indicated. All animal procedures were performed in accordance with protocols approved by the Institutional Animal Care and Usage Committee of New York University School of Medicine.

### Generation of BAC transgenic reporter and CRISPR knockout mice

Tg (Δ+3kb *Rorc(t)*-mCherry) mice were generated as previously described^21^. The following primers were used for generating amplicons for GalK recombineering, and screening for correct insertion and later removal of the GalK cassette: Galk Rec +3kb F: CTGCCTCCCACGTGCTAGGATTGTAATATAGAGCATCAGGCCCTGCTCCACCTGTTGACAATTAATCATCGGCA; Galk Rec +3kb R: ACAGACAGATACCATTCCTTGGGCCTGGCTTCCCTCAGTGGTCCTGGCTGTCAGCACTGTCCTGCTCCTT; +3kb HA F: CTGCCTCCCACGTGCTAGGATTGTAATATAGAGCATCAGGCCCTGCTCCA; +3kb HA R: ACAGACAGATACCATTCCTTGGGCCTGGCTTCCCTCAGTGGTCCTGGCTG; +3kb screen F: CTGCCTCCCACGTGCTAGGAT; +3kb screen R: ACAGACAGATACCATTCCTTGGG. The following primers were used for the recombineering that led to scarless deletion of cis-element: deletion template +3kb HA F: CTGCCTCCCACGTGCTAGGATTGTAATATAGAGCATCAGGCCCTGCTCCACAGCCAGGACCACTGAGGGAAGCCAGGCCCAAGGAATGGTATCTGTCTGT; deletion template +3kb HA R: ACAGACAGATACCATTCCTTGGGCCTGGCTTCCCTCAGTGGTCCTGGCTGTGGAGCAGGGCCTGATGCTCTATATTACAATCCTAGCACGTGGGAGGCAG.

Single guide RNAs (sgRNAs) were designed flanking conserved regions in the +6kb and +7kb region to be deleted using pairs of guides. sgRNAs were cloned into pX458 for use as a PCR template to generate a template for *in vitro* transcription. *In vitro* transcribed sgRNAs and Cas9 mRNA were microinjected into zygotes to generate founder animals that were screened by PCR for deletions. For mice containing expected or interesting deletions, PCR products were TA cloned and Sanger sequenced to determine location of deletion. Founder mice with clean deletions were chosen to breed to C57BL/6 mice to generate lines. The following primers were used to clone sgRNA sequences into BbsI site where lowercase letters represent homology to BbsI: +6/7kb sgRNA-1 F: caccgTTTTCTTTGTGATACCCTTC; +6/7kb sgRNA-1 R: aaacGAAGGGTATCACAAAGAAAAc; +6/7kb sgRNA-2 F: caccGGAGAGACAACTGAAATCGT; +6/7kb sgRNA-2 R: aaacACGATTTCAGTTGTCTCTCC; +6/7kb sgRNA-3 F: caccgTGGACCCAAGTGTTACTGCC; +6/7kb sgRNA-3 R: aaacGGCAGTAACACTTGGGTCCAc.

### Murine antibodies, intracellular staining, and flow cytometry

The following monoclonal antibodies were purchased from Thermo Fisher, BD Biosciences, BioLegend or Abcam: CD3ε (145-2C11), CD4 (RM4-5), CD11b (M1/70), CD11c (N418), CD25 (PC61.5), CD40 (3/23), CD44 (IM7), CD45 (30-F11), CD45.1 (A20), CD45.2 (104), CD62L (MEL-14), CD90.1 (HIS51), CD90.2 (53-2.1), IL-7R (SB/199), CXCR6 (SA051D1 and DANID2), CCR6 (140706), Nkp46 (29A1.4), MHCII I-A/I-E (M5/114.15.2), Ly6G (1A8), Siglec-F (E50-2440), B220 (RA3-6B2), TCR Vα2 (B20.1), TCRβ (H57-597), TCR Vβ5.1/5.2 (MR9-4), TCR Vβ6 (RR4-7), TCRγδ (GL3), IL-22 (IL22JOP), FOXP3 (FJK-16s), RORγt (B2D and Q31-378), GATA3 (TWAJ), T-bet (O4-46), BCL6 (K112-91) and Prdm16 (EPR24315-59). PE-conjugated F(ab’)2-Donkey anti-Rabbit IgG (H+L) (Thermo Fisher) was used as the secondary antibody to detect Prdm16. Anti-mouse CD16/32 (Bio X Cell) was used to block Fc receptors. Live/dead fixable blue (ThermoFisher) was used to exclude dead cells. I-A^b^ OVA_328-337_ tetramers (HAAHAEINEA) were provided by the NIH Tetramer Core Facility.

For labeling OVA-specific T cells, cells were incubated with tetramers for 60 min at 37 °C prior to surface staining. For intracellular staining, cells were stained for surface markers, followed by fixation and permeabilization before intracellular staining according to the manufacturer’s protocol (FOXP3 staining buffer set from Thermo Fisher). For cytokine analysis, cells were stimulated *ex vivo* for 3 h with IL-23 (10 ng/mL; R&D systems) and GolgiStop (BD Biosciences) in complete RPMI-1640 culture medium (RPMI-1640 with 10% FBS, 1% GlutaMAX, 1% penicillin–streptomycin, 10 mM HEPES, and 1 mM sodium pyruvate). Flow cytometric analysis was performed on an LSR II (BD Biosciences) or a Cytek Aurora (Cytek Biosciences) and analyzed using FlowJo software (Tree Star).

### Isolation of lymphocytes

For isolation of cells from lymph nodes and spleens, tissues were mechanically disrupted with the plunger of a 1-ml syringe and passed through 70-μm cell strainers. Bone marrow cells were isolated by flushing the marrow from femur bones using a syringe with RPMI-1640 wash medium (RPMI-1640 with 3% FBS, 10 mM HEPES, 1% GlutaMAX, 1 mM sodium pyruvate, and 1% penicillin–streptomycin). Red blood cells were lysed with ACK lysing buffer (Thermo Fisher). Bronchoalveolar lavage fluid (BALF) cells were collected by flushing the lungs twice with 0.75 ml PBS through a catheter inserted into the trachea.

Lung tissues were minced into small fragments and digested at 37 °C for 45 min with shaking in RPMI-1640 wash medium containing 0.5 mg/ml collagenase D (Sigma) and 0.25 mg/ml DNase I (Sigma). After removal of Peyer’s patches and cecal patches, the intestines were opened longitudinally, cut into 0.5 cm pieces, and washed twice with PBS. Intestine tissues were then shaken in HBSS wash medium (without Ca2^+^ and Mg2^+^, containing 3% FBS and 10 mM HEPES) with 1 mM DTT and 5 mM EDTA at 37 °C for 20 min, repeated twice. After washing with HBSS wash medium (without DTT and EDTA), the tissues were digested in RPMI-1640 wash medium containing 1 mg/ml collagenase D (Sigma), 0.25 mg/ml DNase I (Sigma), and 0.1 U/ml Dispase (Worthington) at 37 °C with shaking for 35 min (small intestines) or 55 min (large intestines). Leukocytes were collected at the interface of a 40%/80% Percoll gradient (GE Healthcare).

### *C. rodentium*-mediated colon inflammation

*C. rodentium* strain DBS100 (ATCC51459; American Type Culture Collection) was grown at 37 °C in LB broth. Mice were inoculated with 0.2 ml of a bacterial suspension (2 × 10^9^ CFU) by oral gavage. Mice were followed for the next 14 days to measure body weight change. Fecal pellets were collected and used to measure *C. rodentium* burden with serial dilutions on MacConkey agar plates.

### *H. hepaticus* culture and oral infection

*H. hepaticus* was provided by J. Fox (MIT). *H. hepaticus* was cultured and administered as previously described^32^. Frozen stock aliquots of *H. hepaticus* were stored in Brucella broth with 20% glycerol and frozen at −80 °C. The bacteria were grown on blood agar plates (TSA with 5% sheep blood, Thermo Fisher). Inoculated plates were placed into a hypoxia chamber (Billups-Rothenberg), and anaerobic gas mixture consisting of 80% nitrogen, 10% hydrogen and 10% carbon dioxide (Airgas) was added to create a micro-aerobic atmosphere, in which the oxygen concentration was 3–5%. The micro-aerobic jars containing bacterial plates were left at 37 °C for 4 days before animal inoculation. For oral infection, *H. hepaticus* was resuspended in Brucella broth by application of a pre-moistened sterile cotton swab applicator tip to the colony surface. 0.2 ml bacterial suspension was administered to each mouse by oral gavage. Mice were inoculated for a second dose after 4 days.

### Adoptive transfer of TCRtg cells

Adoptive transfer of Hh7-2tg CD4^+^ T cells was performed as previously described^32^, with minor modifications. Recipient mice were colonized with *H. hepaticus* by oral gavage 8 days before adoptive transfer. Lymph nodes from CD90.1;Hh7-2 TCR transgenic mice were collected and mechanically disassociated. Naïve Hh7-2tg CD4^+^ T cells were sorted as CD4^+^TCRβ^+^CD44^low/−^CD62L^+^CD25^−^Vβ6^+^CD90.1^+^ (Hh7-2tg), on an Aria II (BD Biosciences). Cells were resuspended in PBS on ice and 100,000 cells were then transferred into congenic isotype-labelled recipient mice by retro-orbital injection. Cells were analyzed 14 days after transfer.

Adoptive transfer of OT-II CD4^+^ T cells was performed as previously described^58^, with minor modifications. Lymph nodes from OT-II;UBC-GFP;*Rag1*^−/−^ mice were collected and mechanically dissociated. Naïve OT-II CD4^+^ T cells were sorted as CD4^+^TCRβ^+^CD44^low/−^CD62L^+^CD25^−^Vα2^+^Vβ5.1/5.2^+^GFP^+^ (OT-II), on the Aria II (BD Biosciences). Cells were resuspended in PBS on ice and 100,000 cells were then transferred into recipient mice by retro-orbital injection. Recipient mice received OVA by oral gavage (50 mg; A5378; Sigma) for 4 consecutive days, followed by drinking water containing OVA (2.5 mg/ml) for an additional 7 days after transfer. Cells from mLN were analyzed 5 days after transfer and cells from intestines were analyzed 12 days after transfer.

### Generation of bone marrow chimeric reconstituted mice

To generate chimeric mice, 4- to 5-week-old wild-type CD45.1 mice were irradiated twice with 500 rads per mouse at an interval of 2–5 h (X-RAD 320 X-Ray Irradiator). One day later, mice were reconstituted with bone marrow cells (3–4 × 10^6^) obtained from wild-type CD45.1/2 mice mixed with either CD45.2 Δ+7kb mutant or littermate control bone marrow cells to achieve a ∼50:50 chimera. Mice were kept for a week on broad spectrum antibiotics (0.8 mg/ml sulfamethoxazole and 0.16 mg/ml trimethoprim), followed by microbiome reconstitution. After at least 8 weeks, tissues were collected for analysis.

### Induction of allergic airway inflammation

Mice were tolerized by oral gavage with 50 mg of OVA for four consecutive days. Six days after the last gavage, the mice were sensitized three times, one week apart, with intraperitoneal injections of 100 μg of OVA mixed with 1 mg of Alum (1:1; Alhydrogel® adjuvant 2%; vac-alu-50; InvivoGen). Seven days after the final sensitization, the mice were challenged intranasally with 40 μg of OVA every other day, for a total of three challenges. Mice were analyzed 24 h after the last intranasal administration.

### OVA/cholera toxin allergy mouse model

To induce tolerance, mice were given 50 mg of OVA by oral gavage once daily for four days. Six days after the final gavage, the mice were sensitized by oral administration of 5 mg of OVA combined with 10 μg of cholera toxin (C8052; Sigma), administered three times at weekly intervals. Seven days after the last sensitization, an intraperitoneal challenge of 200 μg of OVA was administered. Rectal temperature was measured every 15 minutes for a total of 75 minutes after the challenge, using a Type J/K/T thermocouple thermometer (Kent Scientific). Serum was collected the following day to measure anti-OVA IgE and IgG1 levels.

### Histology analysis

Lung tissues were perfused and fixed with 10% buffered formalin phosphate, embedded in paraffin, and sectioned. Lung sections were stained with hematoxylin and eosin and scored for histopathology in a blinded fashion as previously described^59^, with minor modifications. Briefly, lung inflammation was assessed by scoring cellular infiltration around the airways and vessels: 0, no infiltrates; 1, a few inflammatory cells; 2, a ring of inflammatory cells 1-cell-layer deep; 3, a ring of inflammatory cells 2–3 cells deep; 4, a ring of inflammatory cells 4–5 cells deep; and 5, a ring of inflammatory cells greater than 5 cells deep. The inflammation score for each mouse was calculated as the average of the scores from five lung lobes.

Intestine tissues were fixed with 4% paraformaldehyde, embedded in paraffin, and sectioned. Intestine sections were stained with hematoxylin and eosin and scored for histopathology in a blinded fashion as previously described^60^.

### Enzyme-linked immunosorbent assay

Serum levels of total IgE (432404; BioLegend), anti-OVA IgE (439807; BioLegend) and anti-OVA IgG1 (500830; Cayman Chemical) were measured following the manufacturer’s recommendations.

### Bulk ATAC-seq

Samples were prepared as previously described^21,61^. Briefly, 20-50,000 sort-purified cells were pelleted in a fixed rotor centrifuge at 500 g for 5 minutes, washed once with 50 μL of cold PBS buffer. Spun down again at 500 g for 5 min. Cells were gently pipetted to resuspend the cell pellet in 50 μL of cold lysis buffer (10 mM Tris-HCl, pH7.4, 10 mM NaCl, 3 mM MgCl2, 0.1% IGEPAL CA-630) for 10 minutes. Cells were then spun down immediately at 500 g for 10 min and 4 °C after which the supernatant was discarded and proceeded immediately to the Tn5 transposition reaction. Gently pipette to resuspend nuclei in the transposition reaction mix. Incubate the transposition reaction at 37 °C for 30 min. Immediately following transposition, purify using a Qiagen MinElute Kit. Elute transposed DNA in 10 μL Elution Buffer (10 mM Tris buffer, pH 8.0). The transposed nuclei were then amplified using NEBNext High-fidelity 2X PCR master mix for 5 cycles. In order to reduce GC and size bias in PCR, the PCR reaction is monitored using qPCR to stop amplification prior to saturation using a qPCR side reaction. The additional number of cycles needed for the remaining 45 μL PCR reaction is determined as following: (1) Plot linear Rn vs. Cycle (2) Set 5000 RF threshold (3) Calculate the # of cycle that is corresponded to 1/4 of maximum fluorescent intensity. Purify amplified library using Qiagen PCR Cleanup Kit. Elute the purified library in 20 μL Elution Buffer (10mM Tris Buffer, pH 8). The purified libraries were then run on a high sensitivity Tapestation to determine if proper tagmentation was achieved (band pattern, not too much large untagmented DNA or small overtagmented DNA at the top or bottom of gel. Paired-end 50 bp sequences were generated from samples on an Illumina HiSeq2500. Sequences were mapped to the murine genome (mm10) with bowtie2 (2.2.3), filtered based on mapping score (MAPQ > 30, Samtools (0.1.19)), and duplicates removed (Picard).

### Murine mLN collection, cytometric sorting, and scRNA-seq

Live, CD45^+^Lin^−^ APC were sorted (Fig. 2a-d) from the mLN of 3-week-old Δ+7kb mutants (n = 7) and their littermate controls (n = 7), as well as from 17-22 day-old *Rorc(t)*^Δ*CD11c*^ mutants (n = 4) and their littermate controls (n = 4). Lineage markers comprised TCRβ, TCRγδ, B220, and Ly6G. All libraries were prepared using Chromium Next GEM Single Cell 3-prime Kit v3.1 (10x Genomics), following the vendor’s protocol, and sequenced on an Illumina NovaSeq X+ sequencer.

### Murine multiome scRNA-seq/scATAC-seq

Live, CD45^+^Lin^−^MHCII^+^tdTomato^+^ cells, along with one-eighth of the CD45^+^Lin^−^MHCII^+^tdTomato^−^CD11c^+^ cells were sorted (Fig. 2e) from the mLN of 3-week-old *Rorc(t)*^cre^*ROSA26*^wt/lsl-tdTomato^ mice (n = 4) and combined. Nuclear isolation was performed using a demonstrated protocol for single-cell multiome ATAC and gene expression sequencing (CG000365; 10x Genomics). All libraries were prepared using Chromium Next GEM Single Cell Multiome ATAC + Gene Expression Kit (10x Genomics), following the vendor’s protocol, and sequenced on an Illumina NovaSeq X+ sequencer.

### Human mLN collection, cytometric sorting, and scRNA-seq

We coordinated with a transplant surgery operating room to be notified immediately once a deceased (brain dead) donor was identified for life-saving clinical organ harvest. We procured four mesenteric lymph nodes from a 22-year-old patient maintained on life support, with minimal time from surgical resection to digestion of tissue at the benchside, resulting in >92% viable cells. The donor was free of chronic diseases and cancer, and negative for hepatitis B, hepatitis C, and HIV. This study does not qualify as human subjects research, as confirmed by NYU Langone Institutional Review Board, because tissues were obtained from a de-identified deceased individual.

The nodes were transported in Belzer UW Cold Storage Solution, and resected for excess adipose. Digestion buffer comprised RPMI media with 0.2mg/mL Liberase Thermolysin Medium and 2.5mg/mL collagenase D. Each node was pierced with a 31G syringe, injected with digestion buffer until plump, and submerged in the same for 30 minutes at 37 degrees under agitation. Placing onto 100 micron filters, nodes were smashed open with syringe plungers, washing with excess RPMI to maximally recover cells. The suspension was stained with Live/dead eFluor780 (ThermoFisher), CD19 and CD3 (Biolegend HIB19 and UCHT1), and HLA-DR (BD Biosciences L243). Cells were initially gated to enrich viable non-lymphocyte singlets (Fig. 5a). Not knowing MHCII expression levels within human nodes *a priori*, we chose to sort a wide range of HLA-DR^+^ stained cells, omitting only the very bottom quartile. Sorting 61,232 cells ultimately allowed loading of 26,000 across 2 lanes of a 10X microfluidics chip, followed by the same library preparation and sequencing as for murine experiments.

### Computational analysis

RNA-sequencing data were aligned to reference genomes mm10-2020-A (mouse) or GRCh38-2020-A (human) and counted by Cell Ranger (v7.1.0) with default quality control parameters. Each dataset was filtered, removing cells with fewer than 500 detected genes, those with an aberrantly high number of genes (more than 10,000), and those with a high percentage of mitochondrial genes (more than 5%). We computed cell cycle scores for known S-phase and G2/M-phase marker genes, regressing out these variables, along with mitochondrial and ribosomal protein genes.

We employed a completely unsupervised computational workflow that analyzed all cells in aggregate, utilizing the most recent methods^62^ within Seurat version 5.1 (including *sctransform* function) for normalization of gene expression, anchor-based integration (based on 3000 features), and shared nearest neighbor cluster identification. We performed dimensional reduction using a principal component analysis that retained 50 dimensions, and clustering with the Leiden algorithm set to resolution = 1.0. This matched together cells with shared biological states across mutant and control animals, ensuring the final clusters were not driven by effects within either condition. The *Rorc(t)*^Δ*CD11c*^ murine sc-RNA-seq dataset was appended to the Δ+7kb dataset, also clustered in an unsupervised manner, and then analyzed for the same populations as before. All available raw public datasets from Gene Expression Omnibus and Chan Zuckerberg CELLxGENE repositories were filtered with standard quality controls only (no cell types were removed *a priori*), and similarly integrated alongside previous data. The human sc-RNA-seq dataset also underwent a predominantly unsupervised pipeline, but the tDC population was subsequently manually sub-clustered from a juxtaposed cDC2 population.

All sc-ATAC-seq analysis was performed with Signac version 1.14, following a standard workflow^63^. In brief, murine data were annotated with “EnsDb.Mmusculus.v79” and converted to UCSC mm10 annotation where appropriate, and human data were annotated with "EnsDb.Hsapiens.v98" and converted to UCSC hg38 annotation. Density scatter plots established rational thresholds for quality control, filtering data based on transcriptional start site scores, nucleosome banding patterns, features per cell, and mitochondrial genes. Normalization was performed with term frequency-inverse document frequency (TF-IDF). Dimensional reduction was performed with singular value decomposition on the TF-IDF matrix.

DEGs were identified with the Seurat FindMarkers algorithm using the Wilcoxon Rank Sum test, computing the fold change gene expression within each indicated cell type (tDC, cDC1, ILC3) as compared to all other cell types, and this was performed separately within mouse and human datasets. Testing was limited to genes demonstrating Log_2_(Fold Change)=3.2 upregulated genes. The DEGs that defined each cell type according to this threshold, were compared to the other species, as well as compared across indicated cell types, to generate the Venn diagrams in Extended Data Fig. 10b. All 141 DEGs for human tDC are illustrated in the volcano plot in Extended Data Fig. 10c, with the 8 DEGs that are shared by murine tDC specifically annotated.

See “Data and code availability” to access all annotated code in R and Python used to analyze and to generate each Figure panel, all primary raw data, aggregated public raw data, and all post-processed datasets.

### Statistical analysis

Unpaired two-sided t-test, paired two-sided t-test and two-stage step-up method of Benjamini, Krieger, and Yekutieli were performed to compare the results using GraphPad Prism, Version 10 (GraphPad Software). No samples were excluded from analysis. We treated less than 0.05 of P value as significant differences. *P < 0.05, **P < 0.01, and ***P < 0.001. Details regarding number of replicates and representative data can be found in figure legends.

